# A Mathematical Model of Mitochondria in Proximal Tubule and Thick Ascending Limb Cells

**DOI:** 10.1101/2021.12.16.473049

**Authors:** William Bell, Anita T. Layton

**Affiliations:** Department of Applied Mathematics, University of Waterloo, Waterloo, Ontario, Canada; Departments of Applied Mathematics and Biology, Cheriton School of Computer Science, School of Pharmacy, University of Waterloo, Waterloo, Ontario, Canada

## Abstract

Mitochondria are a key player in several kinds of tissue injury, and are even the ultimate cause of certain diseases. In this work we introduce new models of mitochondrial ATP generation in multiple tissues, including liver hepatocytes and the medullary thick ascending limb in the kidney. Using this model, we predict these tissues’ responses to hypoxia, uncoupling, ischemia-reperfusion, and oxidative phosphorylation dysfunction. Our results suggest mechanisms explaining differences in robustness of mitochondrial function across tissues.The medullary thick ascending limb and proximal tubule in the kidney both experience a high metabolic demand, while having lower baseline activity of oxidative phosphorylation relative to the liver. These factors make these tissues susceptible to dysfunction of ComplexIII. A lower baseline oxygen tension observed in the thick ascending limb makes it susceptible to Complex IV. On the other hand, since the liver lacks these risk factors, and has higher baseline rates of glycolysis, it is less susceptible to all kinds of oxidative phosphorylation dysfunction.

## 1 Introduction

Mitochondria are a central organelle for the function of almost all eukaryotic cells, and while our understanding of their significance dates back quite awhile, they’re still capable of surprising us with new mechanisms via which they are important to understanding human diseases [39] [67]. The prevalence of mitochondrial disease for instance was not understood until the sequencing of mitochondrial DNA in the 1990s [67]. The study of the impacts of mitochondrial dynamics in various tissues is a growing area of research *in vivo*, *in vitro*, and *in silico*.

Motivated by the prevalence of chronic kidney disease (CKD) and diabetic nephropathy, the present study focuses on the proximal tubule (PT) and medullary thick ascending limb (mTAL). The large active transport requirements of the PT are a source of stress. The PT accounts for two thirds of sodium reabsorption by the kidney, done mostly via active transport [51]. The work of Edwards et al. [17] that we build on here suggests that the efficiency of cellular respiration in the kidney is reduced by diabetes, suggesting that could be a cause of diabetic nephropathy. Furthermore they suggest that a class of diabetes medications that target the kidney, SGLT-2 inhibitors, could worsen the nephropathic effects of diabetes. The model includes chemical kinetics for pyruvate oxidation, the Krebs cycle, and oxidative phosphorylation, as well as a phenomenological description of the consumption of ATP, an important fuel for the cell. We examine the robustness of the PT in the kidney in light of this large metabolic demand, in response to known stressors upon the PT.

The outer medullar part of the thick ascending limb of the loop of Henle (mTAL) is notable for its contribution to acute kidney injury (AKI) especially brought on by hypoxemia, such as is observed in septic shock [11, 19]. The medulla of the kidney has low tissue oxygen tension but plays a large role in active sodium transport [78], and hypoxemia is more likely to translate into hypoxia in the outer medulla of the kidney due to the inelasticity of its blood supply compared to the renal cortex [19]. Of the nephron segments that are largely in the outer medulla, the mTAL plays the largest role in active reabsorption, and so is at risk due to its high metabolic demand. For this reason, the best way to extend *in silico* methods for studying renal hypoxia to a wider range of cases is to extend them to cover the thick ascending limb in the outer medulla. There is some modelling work on the mTAL, that suggests that the oxygen consumption is more efficient in the mTAL than its active sodium transport activity would suggest, and that this is why the mTAL is robust in the face of its lower baseline oxygen tension [88]. In that paper they hypothesize that this greater efficiency may be due to greater reliance on anaerobic respiration, which we will consider *in silico* using estimates of glycolysis activity in the mTAL. The manner in which the mTAL maintains its active sodium transport activity remains a matter of significant experimental interest with many partial explanations but none that seem conclusive [69].

There are three classes of question that we wish to address in this work. First, the comparative questions: a published *in silico* study on PT mitochondrial function has found that oxygen tension is not rate limiting in the renal cortex for ATP production [17], we compare this to the outer medulla, where lower baseline oxygen tension means that hypoxia may be more extreme. And relatedly, in line with the predictions of Zhang and Edwards [88] that ATP production should be more efficient in the mTAL, we wish to consider the ratios of ATP Synthase mediated ATP generation to Complex IV mediated oxygen consumption, i.e. the P/O ratios (Phosphate/Oxygen ratio), in the PT and mTAL in order to examine their relative efficiencies.

The second kind of question that we hope to answer in this work is about the mTAL itself. The mTAL is known to be subject to greater degrees of hypoxia *in vivo* in the rat under baseline conditions [69], and it is believed that this plays a role in the onset of AKI. Aside from comparative questions mentioned before about its relative robustness, we can directly predict the mitochondrial matrix oxygen tension threshold under which the mTAL will experience hypoxic injury. Evans et al. [19] and Calzavacca et al. [11] report tissue oxygen tensions for various states of hypoxemia and/or shock in the medulla and cortical regions of the kidney in sheep. They find that the kidney might maintain relatively consistent overall oxygenation under ordinary circumstances but during sepsis, renal blood flow was diverted to the cortex over the medulla, and that the outer medulla experienced hypoxia [11]. This accords well with Ma et al. [59] who suggest that septic AKI overall is a disease due to local tissue ischemia. Simulations of the mTAL could address what partial pressures of oxygen tension are required to actually produce pathological tissue hypoxia.

The third kind of question I’ll point to is how the PT and mTAL respectively respond to pathological circumstances, in particular to diabetes, electrolyte or pH imbalance, ischemia-reperfusion cycles, drug-use, and mitochondrial disease, along with the aforementioned hypoxia. Some of this work has been done for the PT although not all; for instance uncoupling and pH imbalance were examined in Edwards et al. [17] for the PT. For this reason, this work is not just useful as an exercise in comparison, but also will break new ground in the study of the PT and the mTAL in diseased states.

Edwards et al. examine the effects of diabetes on mitochondria in the kidney by increasing the rate of mitochondrial uncoupling, or proton leak across the inner membrane of the mitochondria. Friederich-Persson et al. [26] suggests that the deletion of UCP-2 completely prevents uncoupling in the diabetic kidney, and eliminates hypoxia (although increasing oxidative stress). This suggests that we should expect uncoupling to produce significant increases in oxygen consumption in the kidney, which we can test with our model. Since the mTAL is hypothesized by Zhang and Edwards [88] to compensate for low oxygen levels with high efficiency, and proton leak reduces this efficiency, it is possible that diabetic nephropathy could trigger hypoxia in the mTAL. To determine whether diabetes may produce hypoxia, it would be necessary to simulate comparable mitochondrial uncoupling in the PT (which is done already by Edwards et al. [17]) and mTAL.

Mitochondrial disease are understood to have a large role in numerous diseases [67] but have not been captured in previous work using the model adapted here. Key questions in the study of mitochondrial disease revolve around the interactions between dysfunctions of different complexes in the electron transport chain, because it is widely observed that epistatic effects play a large role in mitochondrial disease. A question we wish to address is how ATP production is impacted by different combinations of dysfunctions in the electron transport chain. *In vivo* work on mitochondrial disease mechanism is difficult and suffers from the substantial heterogeneity of mitochondrial disease that makes generalizations difficult [1]. Thus *in silico* work may serve as a welcome supplement to experimental work. For instance, it could identify potential therapeutic targets for therapies in the cases of different combinations of electron transport chain defects. This may be done by seeing which electron transport chain component’s recovery would lead to the greatest recovery of ATP concentration. As a side question, since there is known to be substantial heterogeneity in how distinct tissues are impacted by mitochondrial disease [1], it is worthwhile to see if there are substantial differences between the impact of mitochondrial diseases in the PT and the mTAL.

Ischemia-reperfusion injury has been a major subject of research for those interested in mitochondrial dynamics in the kidney [33]. Transplantation often requires that the donor kidney is not perfused for an extended period of time, but in that case, the kidney may become injured upon reperfusion. More precisely, delayed graft function incidence among kidney transplant recipients is roughly 21% [81] and delayed graft function is understood to be the principle manifestation of ischemia-reperfusion injury. A frequently suggested explanation of this phenomenon is that reperfusion leads to the sudden generation of reactive oxygen species, which, arriving in a pulse, cause significant oxidative damage to the cell. These reactive oxygen species may also be produced before the cell can produce adequate antioxidants to respond to the generation of reactive oxygen species. A helpful starting point for testing these hypotheses is measuring the redox state of the electron transport chain and the proton motive force during reperfusion. Addressing this question would have implications beyond transplantation, because as discussed above, ischemia might be a hallmark of septic AKI as well [59].

## 2 Methods

The present is based on Ref. [17], which treat ATP consumption as independent of the system’s state. This assumption was justified there by the necessity of reabsorbing a large portion of ions from the glomerular filtrate, the necessary level of reabsorption is relatively inelastic. But even if ion reabsorption is necessary, it is still sensitive to ATP levels in the cell, and so cannot be entirely inelastic. For this reason, it is reasonable to choose Michaelis-Menten dynamics for the cell’s ATP consumption (*Q*_ATP_), i.e.

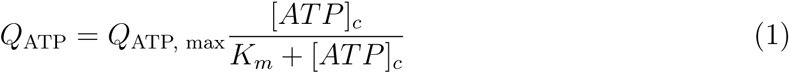

To make the model comparable to the previous model, *Q*_ATP, max_ was chosen so that the ATP consumption was the same at the previous model’s predicted equilibrium. *K_m_* was chosen to equal the *K_m_* of Na-K-ATPase because of its major part in ATP consumption in PT and mTAL cells, its value is taken from Vrbhar, Javorkovam, and Pechanova [83], which models ATP consumption by Na-K-ATPase with Michaelis-Menten dynamics. The *K_m_* is below baseline ATP concentrations at 1.49 mM.

### 2.1 Model Parameters

Most parameters were chosen based on existing experimental estimates. Some were estimated in non-specific kidney tissue when more specific measurements were unavailable. The results are collected in Table 1.

**Table 1:**
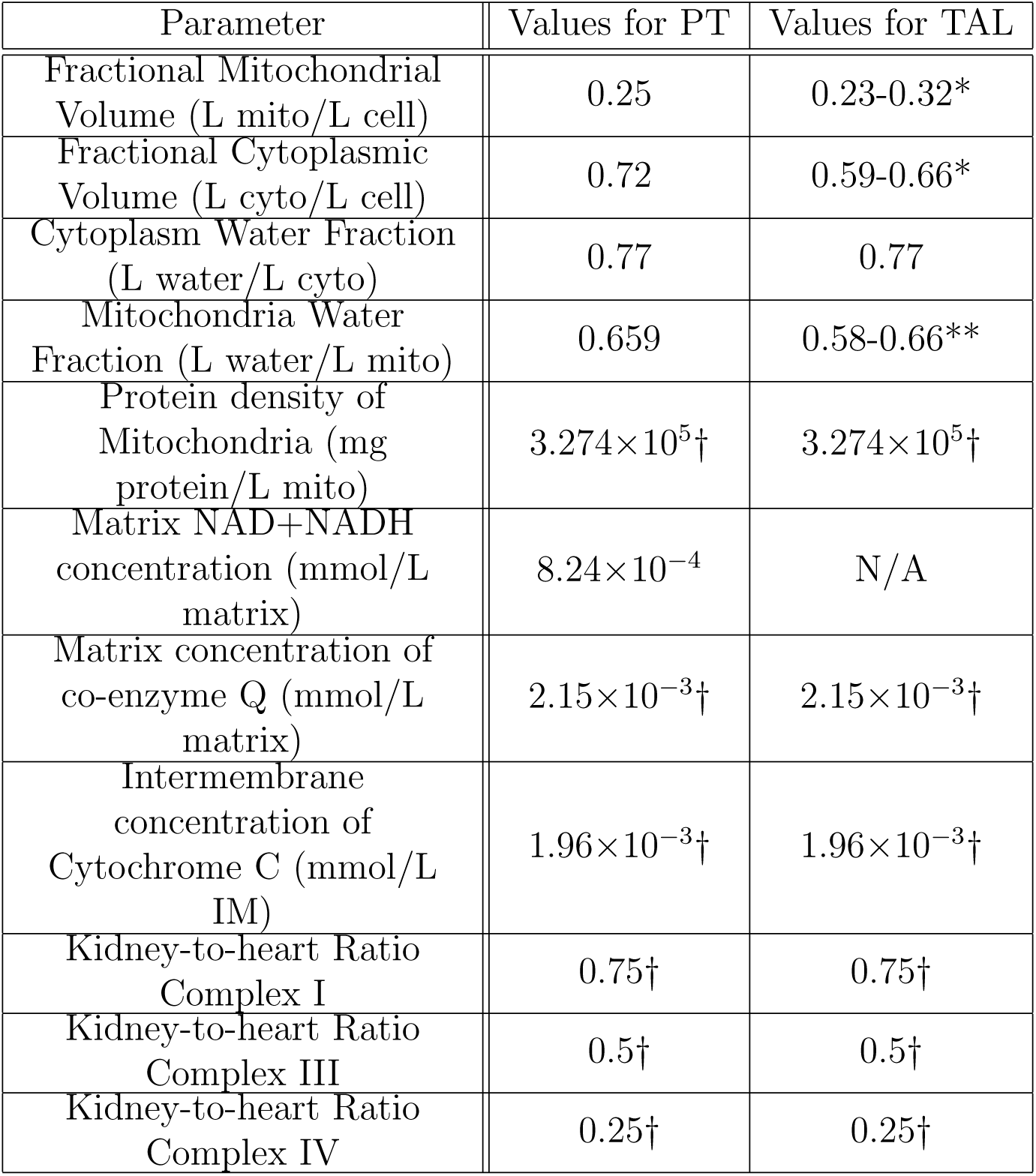
The parameters used in the model where we had to find parameter estimates. The single-starred ranges represent physiological ranges rather than parametric uncertainty. There are large differences between the inner and outer slice of the outer medulla (often much larger than the standard error interval). We use the higher end of the range for the mTAL’s fractional mitochondrial volume and lower end of the range for the fractional cytoplasmic volume, representing the more hypoxic inner slice of the outer medulla. The double-starred ranges represent values chosen to deliberately cover a wide range. The daggers represent values estimated on non-specific kidney tissues. Values marked N/A were fitted rather than chosen based on estimates from the literature. References and further discussion of parameter choices can be found in Section 2.1 and 2.2.

| Parameter | Values for PT | Values for TAL |
| --- | --- | --- |
| Fractional Mitochondrial Volume (L mito/L cell) | 0.25 | 0.23-0.32* |
| Fractional Cytoplasmic Volume (L cyto/L cell) | 0.72 | 0.59-0.66* |
| Cytoplasm Water Fraction (L water/L cyto) | 0.77 | 0.77 |
| Mitochondria Water Fraction (L water/L mito) | 0.659 | 0.58-0.66** |
| Protein density of Mitochondria (mg protein/L mito) | $3.274 \times 10^5 \dagger$ | $3.274 \times 10^5 \dagger$ |
| Matrix NAD+NADH concentration (mmol/L matrix) | $8.24 \times 10^{-4}$ | N/A |
| Matrix concentration of co-enzyme Q (mmol/L matrix) | $2.15 \times 10^{-3} \dagger$ | $2.15 \times 10^{-3} \dagger$ |
| Intermembrane concentration of Cytochrome C (mmol/L IM) | $1.96 \times 10^{-3} \dagger$ | $1.96 \times 10^{-3} \dagger$ |
| Kidney-to-heart Ratio Complex I | $0.75 \dagger$ | $0.75 \dagger$ |
| Kidney-to-heart Ratio Complex III | $0.5 \dagger$ | $0.5 \dagger$ |
| Kidney-to-heart Ratio Complex IV | $0.25 \dagger$ | $0.25 \dagger$ |

The parameter estimates for the proximal tubule were taken from Edwards et al. [17]. The values for both the PT and mTAL of the fractional mitochondrial volume come from Pfaller and Rittinger [64]. Pfaller and Rittinger give estimates for the mTAL in both the inner and outer stripe of the outer medulla. For the former they estimate the mitochondrial volume fraction to be 0.32, and for the latter, 0.23. The same paper gave us cytoplasm volume fraction estimates for both the PT and mTAL [64]. While it was not possible to find cytoplasm water fraction estimates for the mTAL specifically, estimates from across organs, including in a kidney epithelial cell line, have found comparable estimates of roughly 0.77 [30]. This estimate is now a widely shared assumption in studies of cytoplasmic concentrations, including in the kidney [52, 76]. The cytoplasm water fraction may change under hyper/hypotonic conditions or when the cell is growing. Values for mitochondrial water fraction were not forthcoming for the mTAL, and the mitochondrial water fraction did vary significantly between tissues, and so we considered a range chosen to encompass known estimates in other tissues, including the PT [52, 76, 80]. We found that the cell’s state was not sensitive to the mitochondrial water fraction and so we used the same value as for the PT.

The maximal mTAL ATP consumption was chosen so that at typical ATP concentrations (2.5 mM) the cell’s cytoplasmic ATP consumption is comparable to mTAL ATP metabolic demand that is observed from experimental measurements of ATP consumption. Experimentalists have observed 0.64 pmol ATP consumption per second per mm of mTAL [82]. We can calculate the cytoplasmic ATP consumption using the epithelial cell volume per millimeter of tubule of 566 x 10^3^ *µ*m^3^ per mm of tubule [8] and a cytoplasmic volume fraction of 0.60 in the epithelial cell [64]. Using these values, we find an ATP consumption of 1.70 mM/s in the cytoplasm.

The remaining parameters taken from the literature were for non-specific kidney tissues, often studied for the purpose of studying physiological diversity across organ systems. These estimates are used for the PT and the mTAL. The protein density of mitochondria was estimated by Schmitt et al. [71] for the kidney. The ratio of the activity of Complex I between the kidney and heart is taken from Yu et al. [86]. Complex III and Complex IV ratios between the kidney and the heart, as well as matrix concentrations of co-enzyme Q and intermembrane concentrations of Cytochrome C in the kidney are taken from Benard et al. [7].

### 2.2 Model Parameter Fitting

Our model fitting relied on several data sources, namely State 3 respiration, leak respiration, respiratory control ratio, and P/O ratio estimates for the PT [68], P/O ratio and respiratory control ratio for the mTAL [69], and measurements of the response in electrical potential gradient to nigericin in the PT [20]. We use the BFGS algorithm to find an appropriate parameter set, penalizing divergence from the experimental measurements of the above parameters. We compute the difference between the average of the experimental measurements and our predicted value, divided by the standard deviation of the experimental measurements, squared. Then the objective function is the sum of these squares, plus an extra penalty if the P/O ratio was lower in the mTAL than in the PT (using the difference of the PT and mTAL P/O ratios multiplied by a constant of one hundred, chosen to give it comparable weight to other costs). We allowed the adenine nucleotide translocase (*X*_ANT_), potassium-hydrogen antiporter (*X*_KH_), hydrogen & potassium leak permeabilities (*P*_H, leak_ and *P*_K, leak_), and Complex IV oxygen affinity (*k*_O2_) to vary. The most significant change to these parameters was to the potassium-hydrogen antiporter, which is an order of magnitude lower in our model relative to Edwards et al. [17]. The experimental and fitted values from the above fitting may be found in Tables 2 and 3. The NADH/NAD^+^ pool size in the mTAL was also lowered in this model based on measurements of the NADH/NAD^+^ ratio by Hall et al. [32]. The other experimental measurements considered are not sensitive to the NADH/NAD^+^ pool size and so this does not affect our other results.

**Table 2:** Predictions from fitting experimental measurements in isolated mitochondria from the PT in Sciffer et al. [68] and perfused PT in Feldkamp et al [20].

|  | Experimental Values | Predicted Values |
| --- | --- | --- |
| State 3 O<sub>2</sub> Consumption (nmol O<sub>2</sub> mg<sup>-1</sup> s<sup>-1</sup>) | $4.08 \pm 0.6$ | 3.79 |
| Respiratory Control Ratio | $8.5 \pm 0.9$ | 7.26 |
| Leak Respiration (nmol O<sub>2</sub> mg<sup>-1</sup> s<sup>-1</sup>) | $0.37 \pm 0.06$ | 0.52 |
| P/O Ratio | $1.8 \pm 0.1$ | 1.96 |
| Fold Change in dPsi due to Nigericin | 1.05 | 1.06 |

**Table 3:** Predictions from fitting experimental measurements in isolated mitochondria from the mTAL taken from Schiffer, Gustafsson, and Palm [69].

|  | Experimental Values | Predicted Values |
| --- | --- | --- |
| Respiratory Control Ratio | 10.0±1.6 | 8.92 |
| P/O Ratio | 1.93±0.13 | 1.97 |

### 2.3 Simulations

#### 2.3.1 Comparative Questions and Questions about the mTAL

We consider a range of values for several cytosolic ion concentrations, hydrogen & potassium leak permeabilities (*P*_H, leak_ and *P*_K, leak_), and the maximal ATP consumption in the mTAL (*Q*_ATP, max_). This work serves as a counterpart to work already done for the PT [17], which we reproduce with our refitted model. A full list of considered cases for both the PT and mTAL may be found in Tables 5 and 6.

**Table 4:** The parameters that we studied to examine the effects of a drug on mitochondrial respiration. Unless otherwise noted, the parameters are the same as those noted in the appendix to Wu et al. [85]. Where the scale of the effect is not known, a range was considered. Indications for use are taken from Hall, Bass, and Unwin [31]. Dichloroacetate is not known to be nephrotoxic and was instead studied in combination with other drugs.

| Compound | Parameter(s) Affected | Known effect | Indications for Use |
| --- | --- | --- | --- |
| Ifosfamide | $X_{CI}$ | 55% reduction | Cancer |
| Gentamicin | $X_{CI}$, $X_{CIII}$,<br>$X_{CIV}$, $X_{F1}$,<br>$P_{H,leak}$ | OXPHOS reductions like those described above (see Section 2.3.2), $P_{H,leak}$ increased 1.15-, 2-, 5-, or 10-fold | Antibiotic |
| Salicylate | $P_{H,leak}$ | $P_{H,leak}$ increased 1.15-, 2-, 5-, or 10-fold | Analgesia, anti-inflammatory |
| NRTIs/NtRTIs | $X_{CI}$, $X_{CIII}$,<br>$X_{CIV}$, $X_{F1}$ | OXPHOS reductions like those described above (see Section 2.3.2) | Anti-virals |
| Dichloroacetate | $K_{iNADH}$ | A 1.15- or 2-fold increase | Cancer |

**Table 5:** Several key intracellular quantities and their sensitivities to various parameter changes in the mitochondria in the proximal tubule.

|  | O<sub>2</sub><br>Consumption<br>(nmol O<sub>2</sub><br>mg<sup>-1</sup> s<sup>-1</sup>) | ATP<br>Generation<br>(nmol ATP<br>mg<sup>-1</sup> s<sup>-1</sup>) | P/O | Proton<br>Motive Force<br>(PMF, mV) | [ATP]<sub>c</sub><br>(mM) | [ATP]<sub>c</sub>/<br>[ADP]<sub>c</sub> |
| --- | --- | --- | --- | --- | --- | --- |
| **Base Case** |  |  |  |  |  |  |
| P<sub>O2</sub> = 50mmHg<br>pH<sub>c</sub> = 7.20 | 2.55 | 9.86 | 1.93 | 165.16 | 2.27 | 5.31 |
| **P<sub>O2</sub> Variation** |  |  |  |  |  |  |
| P<sub>O2</sub> = 60mmHg | 2.56 | 9.89 | 1.93 | 165.53 | 2.29 | 5.52 |
| P<sub>O2</sub> = 20mmHg | 2.47 | 9.56 | 2.02 | 162.40 | 2.10 | 3.81 |
| P<sub>O2</sub> = 10mmHg | 2.31 | 9.56 | 2.02 | 158.96 | 1.77 | 2.33 |
| **Q<sub>ATP,max</sub> Variation** |  |  |  |  |  |  |
| Q<sub>ATP,max</sub> x 0.75 | 2.04 | 7.58 | 1.86 | 166.14 | 2.44 | 8.48 |
| Q<sub>ATP,max</sub> x 1.25 | 2.93 | 11.53 | 1.97 | 164.56 | 1.92 | 2.86 |
| Q<sub>ATP,max</sub> x 1.50 | 3.12 | 12.39 | 1.99 | 164.30 | 1.52 | 1.70 |
| **pH<sub>c</sub> Variations** |  |  |  |  |  |  |
| pH<sub>c</sub> = 7.40 | 2.31 | 9.19 | 1.99 | 164.72 | 1.91 | 2.79 |
| pH<sub>c</sub> = 7.00 | 2.73 | 10.09 | 1.84 | 164.54 | 2.43 | 8.29 |
| pH<sub>c</sub> = 6.80 | 2.93 | 10.12 | 1.73 | 162.75 | 2.50 | 10.46 |
| **[K<sup>+</sup>]<sub>c</sub> Variations** |  |  |  |  |  |  |
| [K<sup>+</sup>]<sub>c</sub> = 60mM | 2.62 | 10.04 | 1.92 | 167.01 | 2.38 | 7.14 |
| [K<sup>+</sup>]<sub>c</sub> = 140mM | 2.51 | 9.68 | 1.93 | 164.13 | 2.17 | 4.26 |
| **[Mg<sup>2+</sup>]<sub>c</sub> Variations** |  |  |  |  |  |  |
| [Mg<sup>2+</sup>]<sub>c</sub> = 0.2mM | 2.52 | 9.73 | 1.93 | 165.22 | 2.20 | 4.44 |
| [Mg<sup>2+</sup>]<sub>c</sub> = 0.8mM | 2.58 | 10.00 | 1.94 | 165.10 | 2.35 | 6.64 |
| **Variations in H<sup>+</sup>**<br>**and K<sup>+</sup> leak**<br>**permeability** |  |  |  |  |  |  |
| No leaks | 2.26 | 10.18 | 2.25 | 167.75 | 2.38 | 7.04 |
| No H<sup>+</sup> leak | 2.40 | 9.97 | 2.08 | 165.64 | 2.29 | 5.58 |
| No H<sup>+</sup> leak,<br>10x K<sup>+</sup> leak | 1.63 | 4.30 | 1.32 | 157.03 | 0.56 | 0.56 |
| No K<sup>+</sup> leak,<br>10x H<sup>+</sup> leak | 4.14 | 8.97 | 1.08 | 162.76 | 2.13 | 4.02 |
| 10x H<sup>+</sup> leak | 3.88 | 3.59 | 1.15 | 161.77 | 2.05 | 3.50 |
| 10x Leaks | 2.16 | 3.59 | 0.83 | 155.12 | 0.49 | 0.50 |

**Table 6:**
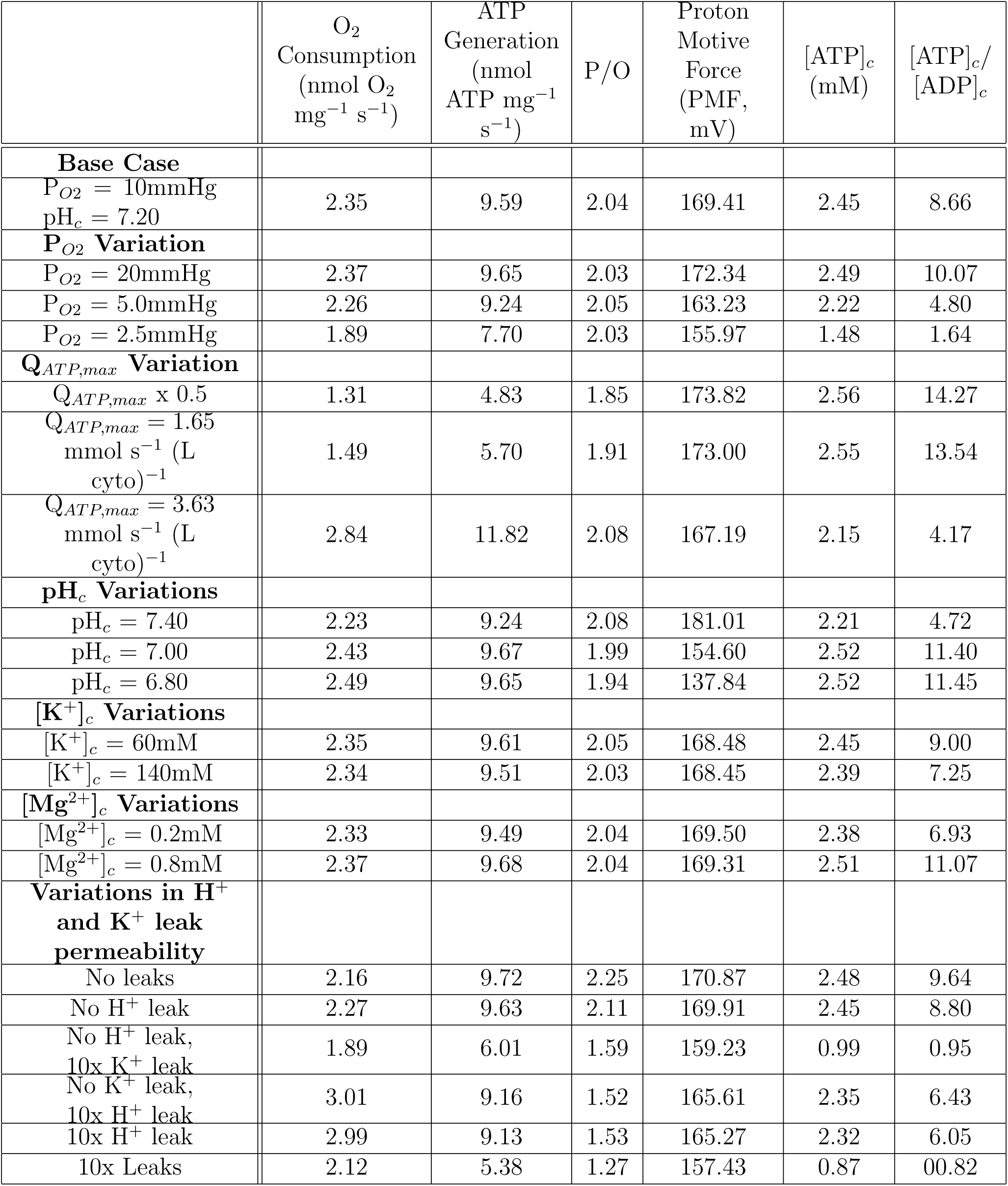
Several key intracellular quantities and their sensitivities to various parameter changes in the mitochondria in the medullary thick ascending limb of the loop of Henle. kidney [24], but on the higher range of in vivo values, as expected for the medullary thick ascending limb [25]. On the other hand, in isolated mitochondria we predict a P/O ratio of 1.77, slightly below the experimentally measured ratio of 1.93 [69]. The concentration of ATP was predicted to be 2.45 mM, the ATP/ADP ratio was predicted to be 8.66, and the proton motive force was predicted to be 169.4 mV. The electrical potential gradient was predicted to be 162.2 mV. Once again these values compare favourably with measurements for the kidney. As previously mentioned previously, typical ranges for the concentration of ATP go from 1.3 to 2.8 mM [54, 15]. Once again our predicted ATP concentration is in the interior of that range. The proton motive force for the mTAL is close to the 170-200 mV range mentioned before [16] and the electrical potential gradient is in the interior of the biologically reasonable 150-180 mV range [9].

|  | O<sub>2</sub><br>Consumption<br>(nmol O<sub>2</sub><br>mg<sup>-1</sup> s<sup>-1</sup>) | ATP<br>Generation<br>(nmol<br>ATP mg<sup>-1</sup><br>s<sup>-1</sup>) | P/O | Proton<br>Motive<br>Force<br>(PMF,<br>mV) | [ATP]<sub>c</sub><br>(mM) | [ATP]<sub>c</sub>/<br>[ADP]<sub>c</sub> |
| --- | --- | --- | --- | --- | --- | --- |
| **Base Case** |  |  |  |  |  |  |
| P<sub>O2</sub> = 10mmHg<br>pH<sub>c</sub> = 7.20 | 2.35 | 9.59 | 2.04 | 169.41 | 2.45 | 8.66 |
| **P<sub>O2</sub> Variation** |  |  |  |  |  |  |
| P<sub>O2</sub> = 20mmHg | 2.37 | 9.65 | 2.03 | 172.34 | 2.49 | 10.07 |
| P<sub>O2</sub> = 5.0mmHg | 2.26 | 9.24 | 2.05 | 163.23 | 2.22 | 4.80 |
| P<sub>O2</sub> = 2.5mmHg | 1.89 | 7.70 | 2.03 | 155.97 | 1.48 | 1.64 |
| **Q<sub>ATP,max</sub> Variation** |  |  |  |  |  |  |
| Q<sub>ATP,max</sub> x 0.5 | 1.31 | 4.83 | 1.85 | 173.82 | 2.56 | 14.27 |
| Q<sub>ATP,max</sub> = 1.65<br>mmol s<sup>-1</sup> (L<br>cyto)<sup>-1</sup> | 1.49 | 5.70 | 1.91 | 173.00 | 2.55 | 13.54 |
| Q<sub>ATP,max</sub> = 3.63<br>mmol s<sup>-1</sup> (L<br>cyto)<sup>-1</sup> | 2.84 | 11.82 | 2.08 | 167.19 | 2.15 | 4.17 |
| **pH<sub>c</sub> Variations** |  |  |  |  |  |  |
| pH<sub>c</sub> = 7.40 | 2.23 | 9.24 | 2.08 | 181.01 | 2.21 | 4.72 |
| pH<sub>c</sub> = 7.00 | 2.43 | 9.67 | 1.99 | 154.60 | 2.52 | 11.40 |
| pH<sub>c</sub> = 6.80 | 2.49 | 9.65 | 1.94 | 137.84 | 2.52 | 11.45 |
| **[K<sup>+</sup>]<sub>c</sub> Variations** |  |  |  |  |  |  |
| [K<sup>+</sup>]<sub>c</sub> = 60mM | 2.35 | 9.61 | 2.05 | 168.48 | 2.45 | 9.00 |
| [K<sup>+</sup>]<sub>c</sub> = 140mM | 2.34 | 9.51 | 2.03 | 168.45 | 2.39 | 7.25 |
| **[Mg<sup>2+</sup>]<sub>c</sub> Variations** |  |  |  |  |  |  |
| [Mg<sup>2+</sup>]<sub>c</sub> = 0.2mM | 2.33 | 9.49 | 2.04 | 169.50 | 2.38 | 6.93 |
| [Mg<sup>2+</sup>]<sub>c</sub> = 0.8mM | 2.37 | 9.68 | 2.04 | 169.31 | 2.51 | 11.07 |
| **Variations in H<sup>+</sup>**<br>**and K<sup>+</sup> leak**<br>**permeability** |  |  |  |  |  |  |
| No leaks | 2.16 | 9.72 | 2.25 | 170.87 | 2.48 | 9.64 |
| No H<sup>+</sup> leak | 2.27 | 9.63 | 2.11 | 169.91 | 2.45 | 8.80 |
| No H<sup>+</sup> leak,<br>10x K<sup>+</sup> leak | 1.89 | 6.01 | 1.59 | 159.23 | 0.99 | 0.95 |
| No K<sup>+</sup> leak,<br>10x H<sup>+</sup> leak | 3.01 | 9.16 | 1.52 | 165.61 | 2.35 | 6.43 |
| 10x H<sup>+</sup> leak | 2.99 | 9.13 | 1.53 | 165.27 | 2.32 | 6.05 |
| 10x Leaks | 2.12 | 5.38 | 1.27 | 157.43 | 0.87 | 00.82 |

#### 2.3.2 Pathological States

Three pathological states are noted above: mitochondrial disease, ischemia-reperfusion injury, and diabetes. Renal Fanconi Syndrome (PT-specific kidney injury) can be caused by mitochondrial disease, often related to a dysfunction of Complex I [3] and especially in pediatric patients. Mitochondrial diseases for the most part impact oxidative phosphorylation, exceptions include impaired mitochondrial biogenesis [2], and so may be represented by reduced activity in the complexes in the inner membrane of the mitochondrion. Specifically we reduce the activity of any combination of Complex I, III, IV, and F_0_F_1_-ATPase (that is, *X*_CI_, *X*_CIII_, *X*_CIV_, and *X*_F1_) by increments of a quarter to as low as a quarter of normal activity.

Ischemia-reperfusion injury is examined here by modelling an extended phase of hypoxia (0 mmHg oxygen tension) followed by reoxygenation, with an adenine nucleotide pool 30% of its typical size, matching the results of Cunningham, Keaveny, and Fitzgerald [14].

Aside from OXPHOS dysfunction, we also consider hypoxia and uncoupling. For hypoxia, we consider reductions in the mitochondrial oxygen tension by tenths to as low as 10% of typical. In the PT the most extreme case considered was an oxygen tension of 5 mmHg and in the mTAL the most extreme case considered was 1 mmHg. For uncoupling (motivated by the study of diabetes, which will be discussed at length in Chapter 5), we consider by default a range of n-fold increases in hydrogen leak activity from twice normal to ten times normal.

#### 2.3.3 Drug Actions

Many cancer drugs and antibiotics directly or indirectly affect cellular respiration. Where their mechanism is well-understood we may estimate the impact of those drugs on cellular respiration, which ultimately helps us better understand how to alleviate the side-effects of those drugs. In particular, drug-induced Fanconi syndrome is associated with mitochondrial dysfunction on both biochemical and histological evidence [31]. We systematize the understanding of these drug actions by a sensitivity analysis of the relevant parameters.

Here we discuss where known effects of different drugs, as recorded in Table 4, were found. The value for how ifosfamide affects Complex I activity is taken from Nissim et al. [62]. The mechanism for gentamicin is through an uncoupling effect and inhibition of magnesium-dependent components of oxidative phosphorylation, which is chiefly via *F*_0_*F*_1_- ATPase [65, 84]. Salicylate, a biproduct of aspirin, is a known uncoupler [60]. Nucleoside(/Nucleotide) reverse transcriptase inhibitors (NRTIs/NtRTIs) like tenofovir impact mitochondrial DNA replication by targetting mitochondria-specific DNA polymerases, this primarily causes general OXPHOS dysfunction like in mitochondrial disease [56].

Dichloroacetate is a cancer drug that is used to switch cells towards aerobic over anaerobic respiration, contrary to the Warburg effect which is cancer’s natural tendency to prefer anaerobic respiration even in the presence of oxygen [77]. Dichloroacetate accomplishes this by preventing pyruvate dehydrogenase kinase from downregulating pyruvate dehydrogenase activity, a key step in pyruvate oxidation. Recent work has examined the possibility of dichloroacetate having renal protective effects in combination with mitotoxic agents such as the uncoupler 2,4-dinitrophenol, an illicit “slimming” drug, with encouraging results [28] and with cisplatin, a cancer drug known to cause Fanconi syndrome [29, 31]. We simulate the combination of dichloroacetate with the previously mentioned drugs to predict whether it may protect against their nephrotoxic effects.

## 3 Results

### 3.1 Baseline Results for the Proximal Tubule

Table 5 includes some baseline numbers and the sensitivity to changes in multiple quantities, both parameters and states for the PT. We consider several predicted values related to both fluxes in the mitochondria and states of the system. Oxygen consumption was predicted to be 2.55 nmol *O*_2_ mg*^−^*^1^ s*^−^*^1^, and at the same time there was ATP generated at a rate of 9.86 nmol (mg prot)*^−^*^1^ s*^−^*^1^ by ATP Synthase. Together these numbers underlie the P/O ratio *in vivo* which was found to be 1.93. Known estimates of the P/O ratio for the kidney range from 1.7-2 *in vivo* [25] to 2.5 in the perfused kidney [24], while these are for the whole kidney, they include our estimate of 1.93. Further the concentration of ATP was predicted to be 2.27 mM, the ATP/ADP ratio was found to be 5.31, and the proton motive force across the inner membrane was predicted to be 165 mV, which together give an account of the cell’s energetic state. These are key variables for which we have experimental results to which to compare, the typical range of proton motive force is from 170 to 200 mV [16]. The mitochondrial electrical potential gradient was predicted to be 155 mV, in the interior of known range from 150 to 180 mV [9]. Typical measured ranges for the concentration of ATP vary widely from 1.3 to 2.8 mM [15, 54]. The ATP/ADP ratio has been measured to be roughly 9 in mice renal cortex [87], however Zager notes that the ratio is extremely variable. Results from others for the rat renal cortex indicate the ratio may be as low as 3, based on independent calculation of the ATP/ADP ratio [34]. Thus a value of 5.31 is within a plausible range for the ATP/ADP ratio. We see that the proton motive force is close to our predicted value, the predicted ATP concentration is in the middle of the possible range, and measured ATP/ADP ratios and electrical potential gradient appear to allow for our prediction. These results for the PT are relatively similar to the values found by Edwards et al. [17]. The proton motive force and the ATP/ADP ratio predictions differ most from the predictions of Edwards et al. (Edwards et al. predict a proton motive force of 165.16 and an ATP/ADP ratio of 10.94), but the latter is very variable as previously mentioned. The latter was found to be sensitive to the parameter variations considered in Table 5.

Across all simulations considered, the P/O ratio or efficiency of the cell was most sensitive to hydrogen and potassium leaks. These sensitivity cases were significant because the mitochondrion’s ion leak activity was highly uncertain. Greater leak decreased efficiency, a tenfold increase in both leaks reduced the P/O ratio to 0.83 (see the last row of Table 5), the lowest predicted for the PT, primarily via decreased ATP generation. In the absence of any leak, the P/O ratio was 2.25 (see the first row of the last block of Table 5).

The greatest factors controlling oxygen consumption were maximal ATP consumption (*Q_ATP,max_*) and hydrogen leak (the third and last blocks in Table 5 include these numbers). The greatest oxygen consumption predicted was 4.14 nmol mg*^−^*^1^ s*^−^*^1^ when potassium leak was zero and hydrogen leak was increased tenfold. At 3.12 nmol per milligram per second, we see that a 1.5 fold increase maximal ATP consumption also increases the oxygen consumption substantially.

The proton motive force was insensitive to most changes, aside from leak permeability changes. The largest proton motive force was found to be 168 mV in the absence of leak, and when both potassium and hydrogen leak were increased tenfold the proton motive force was found to be 155 mV.

The cytosolic potassium and potassium leak (see the fifth and last blocks of Table 5) are impactful on ATP generation, ATP concentration, and the ATP/ADP ratio as well. The lowest ATP generation is 3.59 nmol ATP per milligram per second, the lowest cytosolic ATP concentration is 0.49 mM, and the lowest ATP/ADP ratio is 0.50, all in the case where potassium and hydrogen leaks are both increased tenfold. Results for tenfold increased potassium leak alone are comparable. The maximal ATP generation grows with maximal ATP consumption, for a 1.5-fold increase in maximal ATP consumption, the ATP generation grows 1.3-fold, and for 75% of the typical ATP generation, we see ATP generation go down by roughly the same amount.

### 3.2 Baseline Results for the medullary Thick Ascending Limb of the Loop of Henle

Table 6 includes some key baseline predictions and the model sensitivity to changes in multiple quantities, both parameters and states for the mTAL, analogous to those for the PT above. The oxygen consumption was predicted to be 2.35 nmol *O*_2_ mg*^−^*^1^ s*^−^*^1^, compared to ATP generated at a rate of 9.59 nmol (mg prot)*^−^*^1^ s*^−^*^1^ (or 1.0 mmol (L cell)*^−^*^1^ s*^−^*^1^) by ATP Synthase. Together these numbers get us the P/O ratio *in vivo* which was found to be 2.05. This is slightly below whole kidney P/O measurements noted before for the perfused

The P/O ratio is most sensitive to ATP consumption and the leak permeabilities. At the low end of simulated ranges for ATP consumption, the P/O ratio is 1.85, whereas at the high end of the considered range for ATP consumption, the P/O ratio was predicted to be 2.08. In response to changes in leak permeability we also see large changes in the P/O ratio. In the absence of hydrogen or potassium leak, the P/O ratio is 2.25, and in the worst case scenarios for the P/O ratio, when hydrogen leak was 10 times normal and the potassium leak was 10 times normal, the P/O ratio was 1.27.

In the mTAL, the proton motive force is sensitive to hypoxia, at 2.5 mmHg oxygen tension, we see a proton motive force of 156 mV. The mTAL’s proton motive force is more sensitive to hypoxia because the degree of hypoxia considered leads to much lower electron transport flux for the mTAL than the hypoxia considered for the PT (see the oxygen consumption in the third block in Tables 5 and 6, which is a proxy for electron transport activity). We also see a strong sensitivity of the proton motive force to the cytosolic pH, for a slightly acidic cytosolic pH we see a proton motive force of 138 mV. Predictably, and like for the PT, we also observe that the proton motive force is sensitive to hydrogen and potassium leak, particularly the potassium leak. Under tenfold increased hydrogen and potassium leak capacity, we predict a proton motive force of 157 mV.

### 3.3 Local Sensitivity Analysis for the Proximal Tubule and Thick Ascending Limb

We consider the change in state variables in the PT under a change in each parameter of our model. The full local sensitivity analysis is reported in Figure 1. Here we focus on several key state variables and the parameters that are most impactful upon these state variables. In Figure 3 we show the derivative of each state variable against each of the most significant parameters, calculated using a central difference scheme with Δ*p* = 0.01*p* where *p* is the size of the parameter. The derivative is normalized relative to the baseline size of each state variable and the size of each parameter. We see that the concentrations of NADH and reduced coenzyme Q (QH_2_) are most sensitive to the parameter changes we considered. The cytosolic ATP concentration is most sensitive to the maximal ATP consumption. The concentration of NADH in the cell is determined heavily by alphaketogluterate dehydrogenase, which is positively associated with increased NADH content, and pyruvate dehydrogenase, which is negatively associated with NADH content. Greater alphaketogluterate dehydrogenase activity allows for more production of NADH because in the proximal tubule it is predicted to be a limiting step for the TCA cycle. Alphaketogluterate concentrations are higher in our model than any other TCA cycle intermediate, and the concentration of the product of alphaketogluterate dehydrogenase, succinyl-CoA, is lower than any TCA cycle intermediate. Pyruvate dehydrogenase on the other hand reduces NADH concentrations because it competes with alphaketogluterate dehydrogenase for coenzyme A as a substrate. Pyruvate dehydrogenase uses coenzyme A to produce acetyl-CoA whereas alphaketogluterate uses coenzyme A to produce succinyl-CoA. In the full sensitivity plot, Figure 1, we see that the TCA cycle intermediates from citrate to alphaketogluterate are present in greater concentrations when you increase pyruvate dehydrogenase activity, and intermediates downstream of alphaketogluterate dehydrogenase are present in lower quantities. Supplementary simulations found that for 1% or 10% increases in the activity of pyruvate dehydrogenase, we observe a consistent alphaketogluterate dehydrogenase flux, despite larger concentrations of alphaketogluterate. This observation further supports this proposed mechanism, since more alphaketogluterate is necessary to produce the same enzyme flux. Aside from the above mechanism and its downstream effects, we see unsurprisingly that the concentration of reduced coenzyme Q and reduced cytochrome C continues to increase with increased total quantities of coenzyme Q and cytochrome C. We also see that the passing of electrons from coenzyme Q to cytochrome C appears to be limited by the available cytochrome C, since the reduction state of coenzyme Q appears to be highly sensitive to the total cytochrome C concentration.

**Figure 1:**
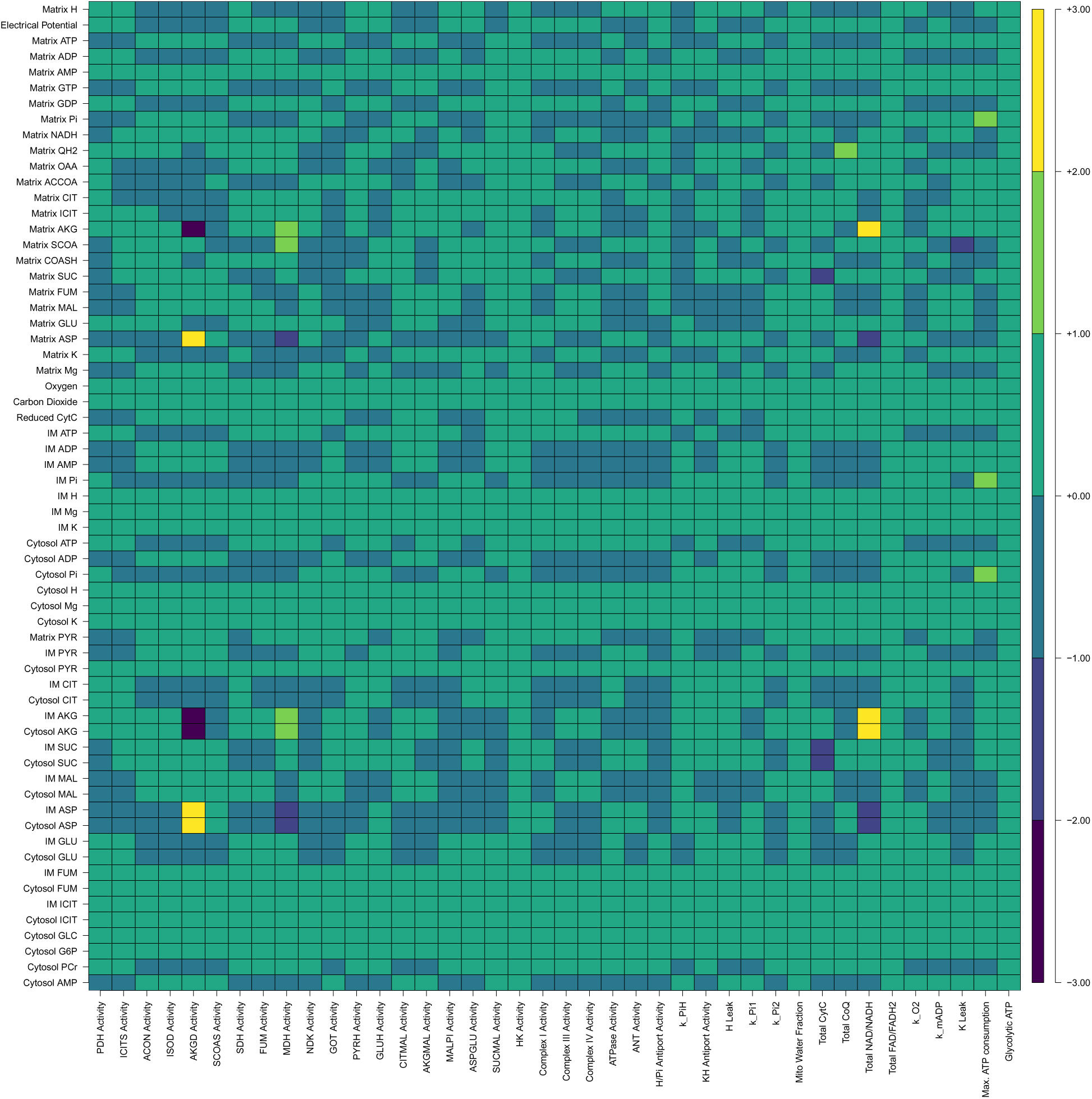
The full set of local sensitivities in the PT, for more information see section 3.3.

In the mTAL we find several differences and an overall greater robustness to small perturbations of key parameters. The full local sensitivity analysis is reported in Figure 2. In Figure 3 we show the sensitivity calculated in the same manner as for the PT. At the same threshold for a parameter to be significant, two more parameters are significantly sensitive in the PT than are sensitive in the mTAL. The concentrations of NADH, reduced coenzyme Q (QH_2_), and reduced cytochrome C are particularly sensitive state variables for the mTAL. The most significant difference between the PT and mTAL is that in the mTAL, NADH concentrations are less sensitive to pyruvate dehydrogenase and alphaketogluterate dehydrogenase activities. Instead we see in the mTAL that the pooled concentration of NADH and NAD^+^ is by far the most important factor impacting the concentration of NADH. This suggests that whereas in the PT there are limited amounts of coenzyme A and this limits the amount available to alphaketogluterate dehydrogenase (as illustrated by the sensitivity to pyruvate dehydrogenase activity), instead NAD^+^ is in short supply in the mTAL, and so the way to provide more NADH to the cell is to increase the size of the NADH/NAD^+^ pool. Given the higher NADH to NAD^+^ ratio in the mTAL and its relative hypoxia (which can leave the electron transport chain in a more reduced state [35]), coenzyme Q is also in high demand, leading to a high sensitivity of multiple states to the pooled coenzyme Q concentration.

**Figure 2:**
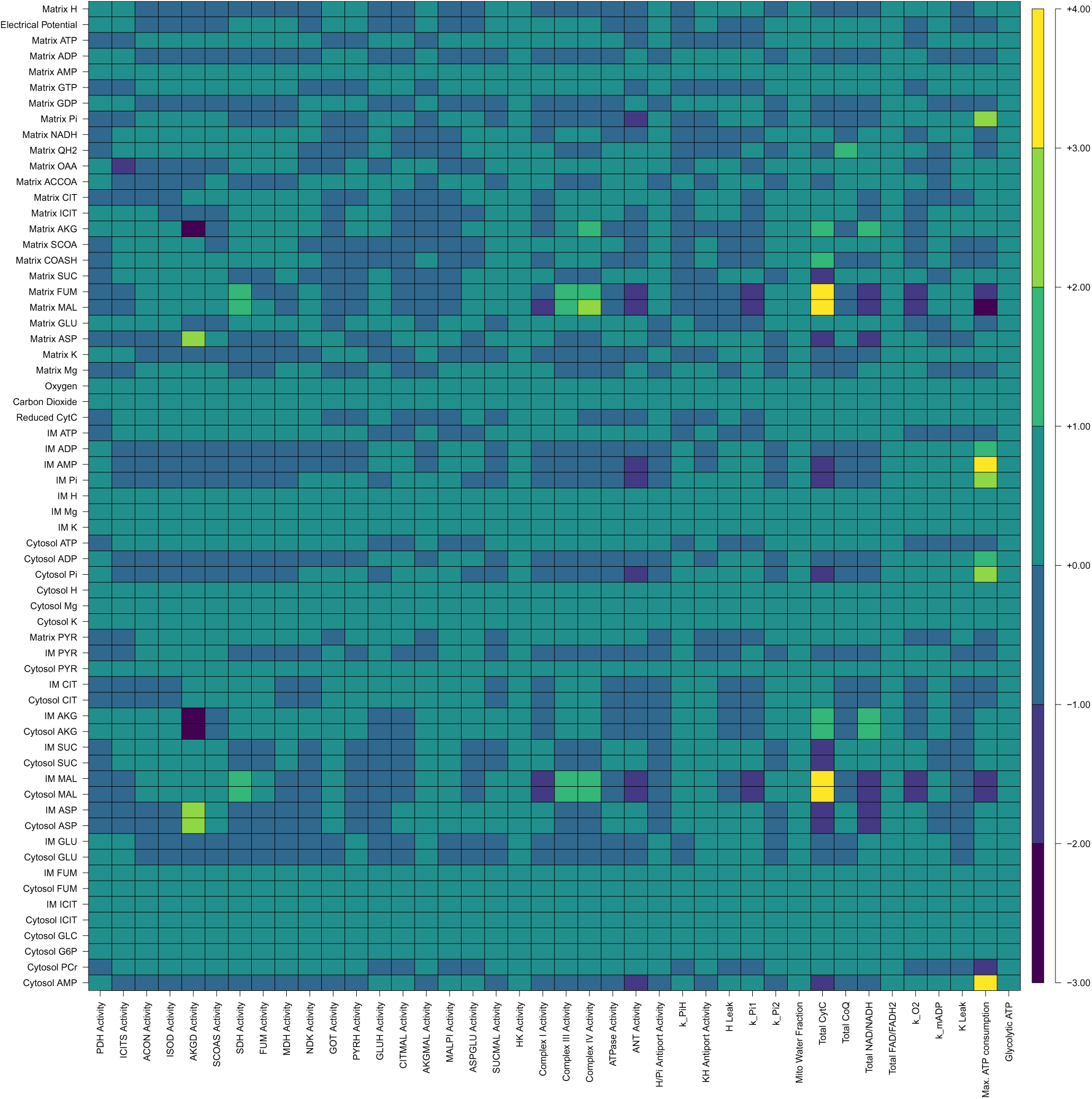
The full set of local sensitivities in the mTAL, for more information see section 3.3.

**Figure 3:**
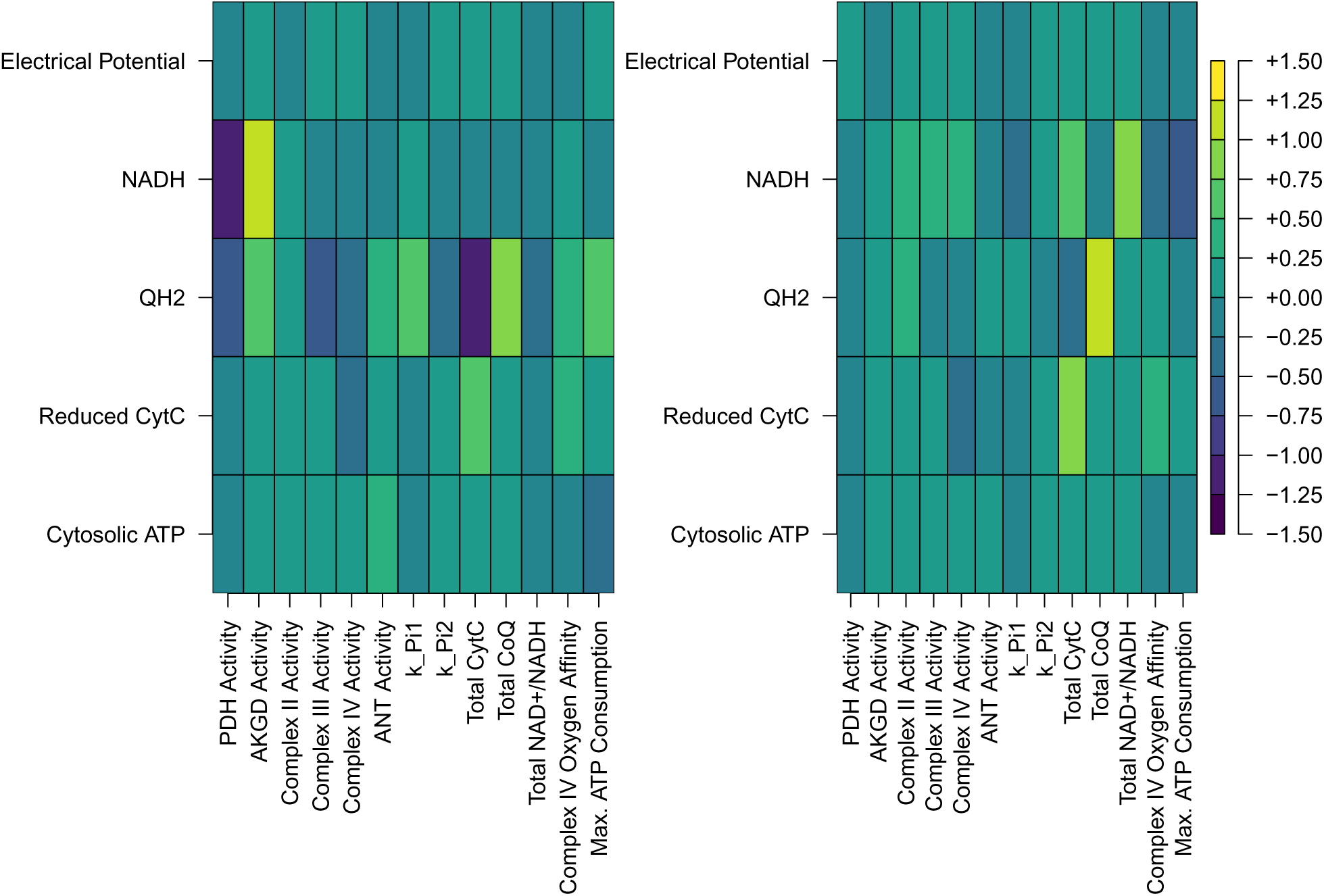
Local sensitivities of several important state variables in the PT (left panel) and mTAL (right panel) relative to certain parameters significant to either the PT or mTAL. The results are calculated as described in section 3.3. CytC stands for cytochrome C, reduced cytochrome C refers to cytochrome C that has been donated an electron by Complex III. AKGD refers to alphaketogluterate dehydrogenase, PDH refers to pyruvate dehydrogenase, ANT refers to adenine nucleotide translocase, and QH_2_ is the reduced form of coenzyme Q.

### 3.4 Global Sensitivity Analysis of the Model

For our global sensitivity analysis we used the Sobol method for uniformly exploring a high-dimensional parameter space and describing the observed variation in the state space as a function of the parameters. For local estimates, interactions are negligible, but in a global sensitivity analysis interactions may be very important. For this reason we report two sensitivity estimates, the first Sobol indices, which do not account for interactions and the total Sobol indices, which account for two-way interactions. First Sobol indices are non-negative numbers between zero and one, whereas total Sobol indices may be greater than one. They represent the proportion of variance explained by a particular parameter in the case of the first Sobol index, and in the case of the total Sobol index, the variance explained by the parameter and the pair-wise interactions associated with it. In Figure 6 we present the first Sobol indices for several important state variables, and in Figure 7 we present the total Sobol indices for those state variables. We see that the first Sobol index results share some features with the results above. We see that like for the local sensitivity analysis of the PT, pyruvate dehydrogenase features prominently, its importance is discussed above in our local sensitivity analysis. Succinyl-CoA synthetase *in vivo* catalyzes the conversion of succinyl-CoA into succinate and coenzyme A. This may increase the concentration of NADH by increasing the availability of coenzyme A, which in the PT and mTAL is found in lower concentrations than acetyl-CoA and succinyl-CoA. This could explain its effects on NADH concentrations in the mitochondrion. The next-most important parameter for influencing the state variables is the maximal ATP consumption, which plays a large role in determining the reduced cytochrome C concentration. ATP consumption frees ADP for phosphorylation. This relieves the proton gradient across the inner membrane. Complex IV requires protons in the mitochondrial matrix in order to oxidize cytochrome C. When we consider pairwise interactions via the total Sobol index we see differences in which parameters stand out. Pyruvate dehydrogenase and succinyl-CoA synthetase still impact coenzyme Q and NADH concentrations. Most notably the activities of adenine nucleotide translocase, alphaketogluterate dehydrogenase, and potassium leak are all significant when we consider interactions. Aside from this the results are fairly similar, and we don’t see large changes in the influence of the most impactful parameters on the important state variables.

**Figure 4:**
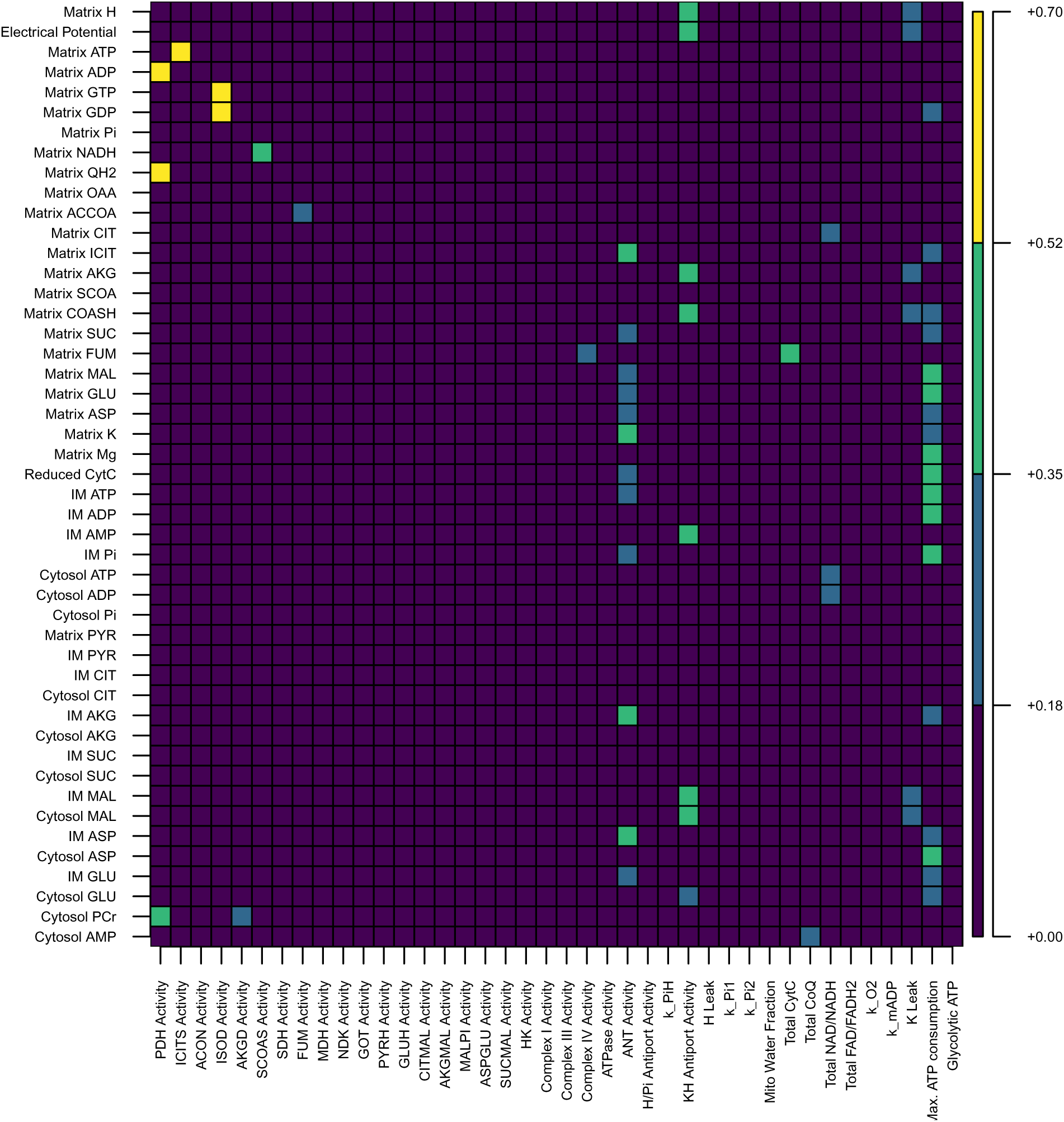
The sensitivity of all state variables to the parameters as calculated according to the Sobol method, without interactions.

**Figure 5:**
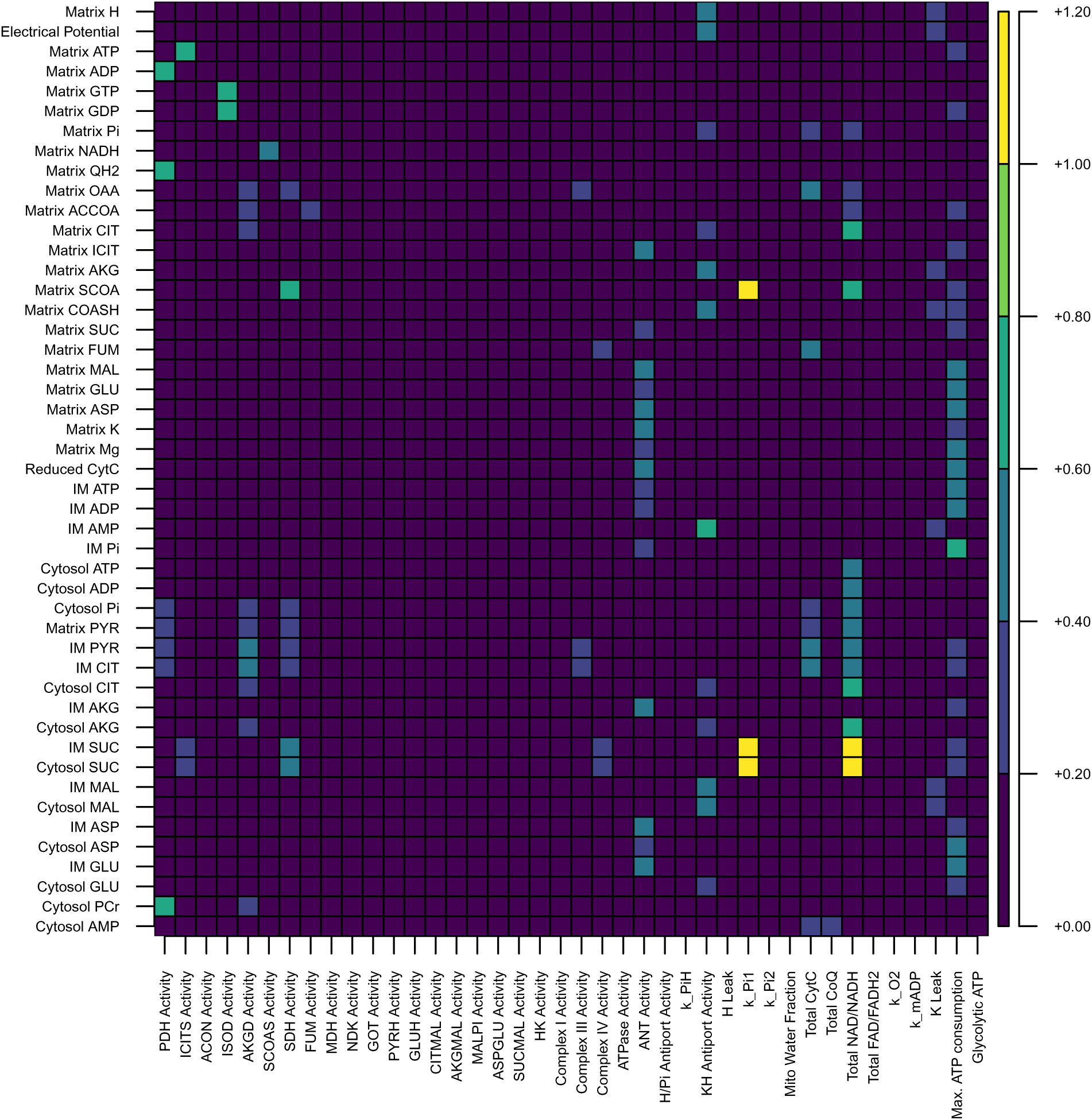
The sensitivity of all state variables to the parameters as calculated according to the Sobol method, taking into account two-way interactions.

**Figure 6:**
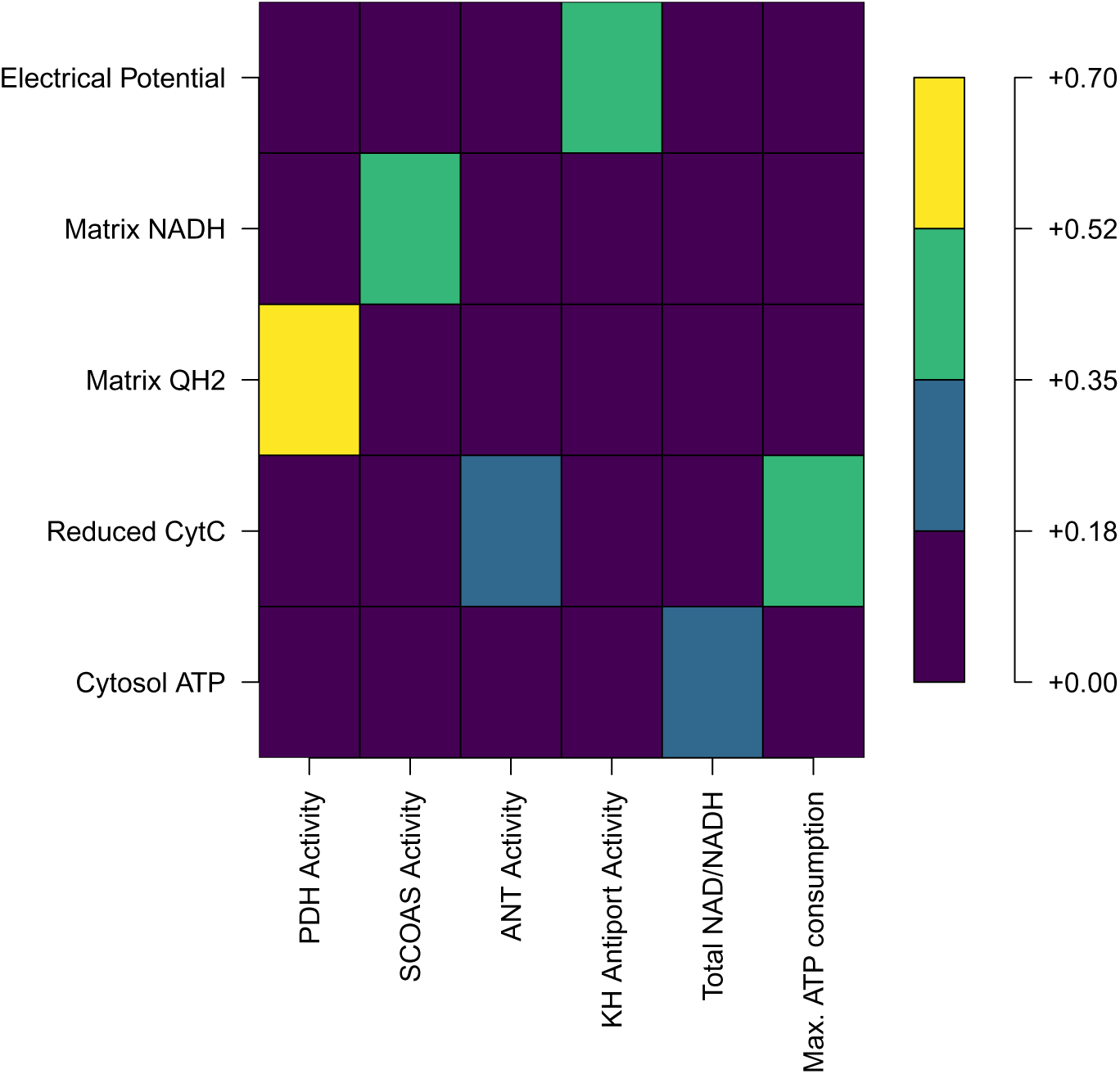
The sensitivity of important state variables to the most important parameters as calculated according to the Sobol method, not including interactions. CytC stands for cytochrome C, reduced cytochrome C refers to cytochrome C that has been donated an electron by Complex III.

**Figure 7:**
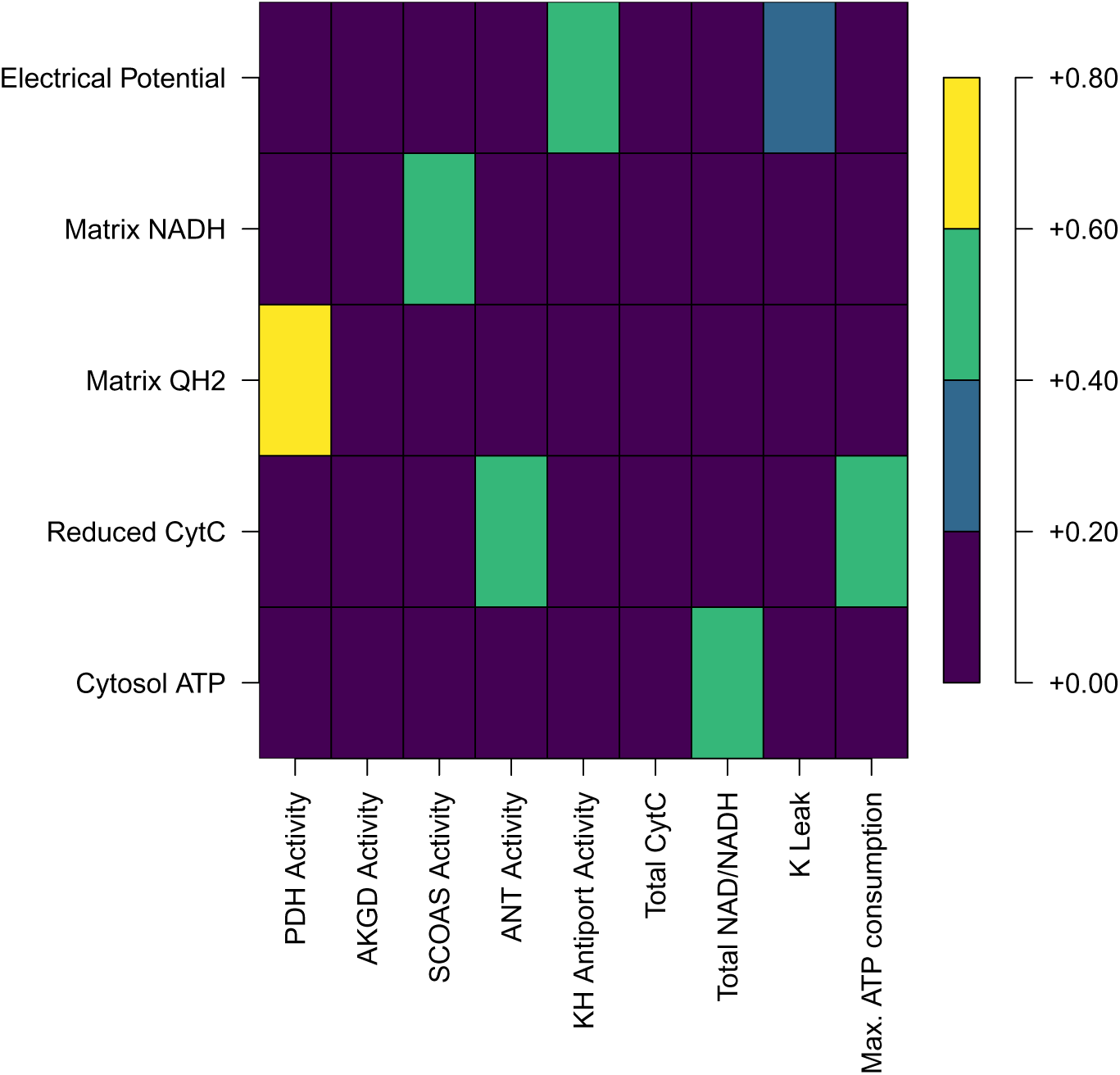
The sensitivity of important state variables to the most important parameters as calculated according to the Sobol method, taking into account two-way interactions. CytC stands for cytochrome C, reduced cytochrome C refers to cytochrome C that has been donated an electron by Complex III.

### 3.5 Response to Nigericin

Nigericin is a potassium-hydrogen antiporter [10] that in the quantities administered experimentally is known to remove entirely the proton gradient across the inner membrane of the mitochondria [43]. Feldkamp et al. [20] take advantage of this to measure the ΔpH, upon adding nigericin, the electrical potential gradient increases by the same amount as the ΔpH decreases. For this reason the addition of nigericin was modelled as an increase in potassium-hydrogen antiport activity, enough to force the ΔpH to zero, we use a twenty-fold increase in antiport activity. We see that this saturates the corresponding increase in electrical potential gradient in Figure 8 (the baseline curve is green). The impact of nigericin on the electrical potential gradient, which goes up as hydrogen gradient is turned into electrical potential gradient, is predicted to go from 160 mV to 168 mV. While the results in Feldkamp et al. [20] are not numerically recorded and are no longer obtainable (Feldkamp, personal communication), based on careful measurement of Figure 4 found in Feldkamp et al. [20], a 5% increase in potential gradient was found to be the approximate effect of nigericin on the electrical potential gradient, after model fitting using this estimate, we see a 6% increase in our model.

**Figure 8:**
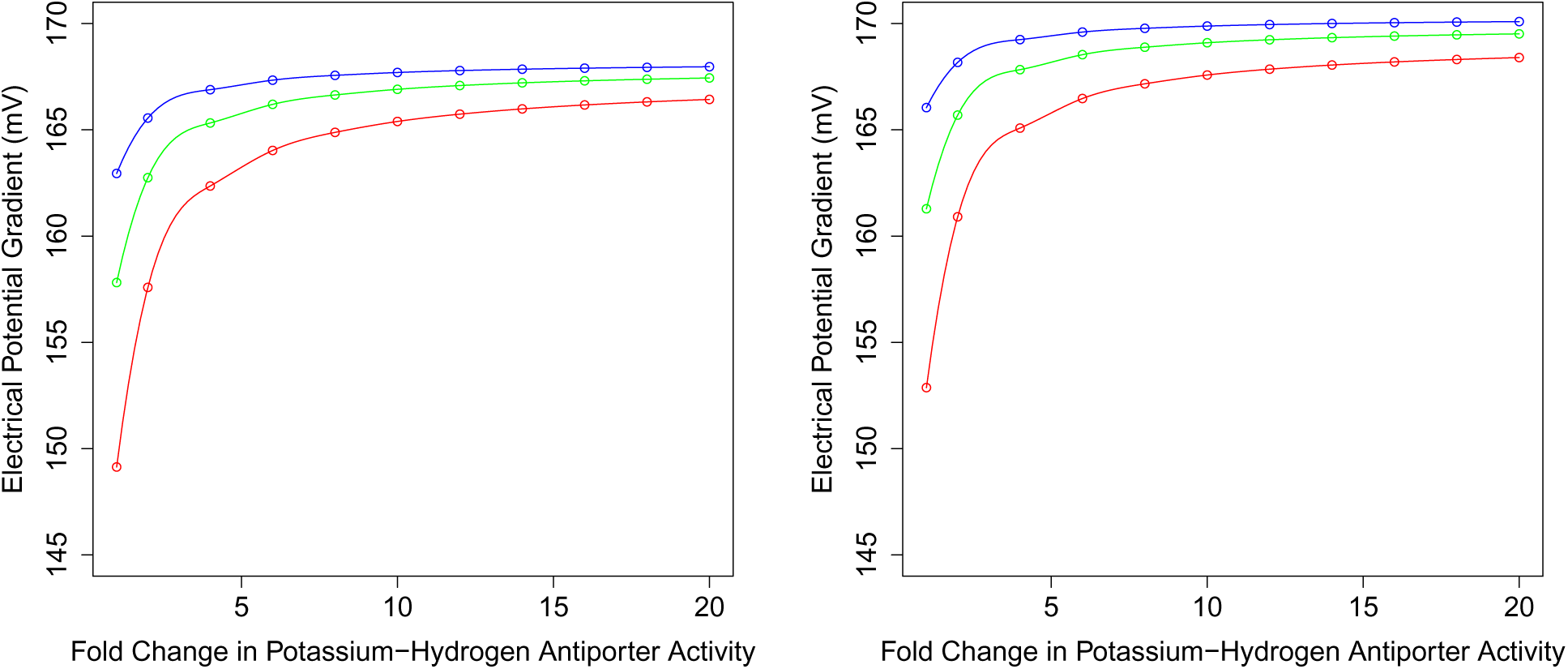
The associated increases in electrical potential gradient in the PT (left) and mTAL (right) for changes in the potassium-hydrogen antiporter activity. The green curve is for baseline potassium leak activity, red is for doubled potassium leak, and blue is for halved potassium leak. The green curve shows a roughly 6% increase in electrical potential gradient from left to right. The curve is a standard cubic spline.

For increased potassium leak, we predict a lower baseline electrical potential gradient. The electrical potential gradient increases under larger potassium-hydrogen antiporter activity, and it increases more (11%) for higher levels of potassium leak (in Figure 8 we see a electrical potential gradient curve in antiporter activity for doubled potassium leak activity in red, and for halved potassium leak in blue). With a large potassium-hydrogen antiporter activity, the electrical potential gradient is less sensitive to potassium leak. This suggests nigericin is more impactful when the potassium leak activity is greater.

Unlike the PT, we did not have experimental data for the use of nigericin in the mTAL. However, using the same procedure as for the PT (a twentyfold increase in potassium-hydrogen antiport activity), we predict the electrical potential gradient increases from 161 mV to 170 mV (6%) in the presence of nigericin for baseline potassium leak activity.

Nigericin is frequently used in studies on mitochondria [20, 41, 89] and so it is valuable that the model correctly captures the response to nigericin. Under previous versions of this model [17], the model did not correctly predict the response found in Feldkamp et al. [20] because the predicted ΔpH was too low. As discussed above in Section 2.1, we use a lower potassium-hydrogen antiporter activity. The potassium-hydrogen antiporter impacts the robustness of the electrical potential gradient to ion leakage.

### 3.6 Uncoupling Effects in Proximal Tubule Mitochondria

Uncoupling or increased hydrogen leak is another mechanism that impacts cell function by reducing the efficiency of respiration. Uncoupling may be caused by diabetes in which there’s an upregulation of UCP-2 [26], or by salicylate, also known as aspirin, a known uncoupler and occasional cause of Fanconi Syndrome [31]. The cases considered here are of uncoupling without any other effects. These cases give a baseline expectation for what might happen during uncoupling. The exact simulation procedure used is outlined in Section 2.3.2. We find that the effect on cytosolic ATP concentration (~0.2 mM in the worst case), proton motive force (≤3 mV in the worst case) and electrical potential gradient (≤1 mV in the worst case) was minimal. The worst case hydrogen leak (tenfold increased hydrogen leak activity) considered is included in Table 5. The minimal effects of hydrogen leak on ATP generation and mitochondrial polarization are explained by compensatory increases in the flux through Complex I, III, and IV in our model. These electron transport chain components pump protons across the inner membrane into the intermembrane space, and so with increased hydrogen leak, they have more substrate to do their work. This is compatible with the hypothesis that uncoupling can perform an adaptive function by providing an outlet for over-reduced oxidative phosphorylation complexes [26]. This would not be possible if electrical potential gradient was significantly lost during mild uncoupling [85]. As noted in Section 3.1, under increased hydrogen leak, the oxygen consumption by the proximal tubule may be as much as 52% higher, this could be enough to trigger hypoxia, possibly causing progression of diabetic nephropathy [26]. This is an unfortunate consequence of the previously mentioned increases in Complex I, III, and IV activity. The oxygen consumption included in Table 5 is directly proportional to Complex IV activity.

### 3.7 Uncoupling Effects in the medullary Thick Ascending Limb Mitochondria

Uncoupling can also happen in the mTAL, and the effects were studied similarly. The effects on the cytosolic ATP concentration (*<*0.1 mM in the worst case), proton motive force (~11 mV in the worst case), and electrical potential gradient (~5 mV in the worst case) were minimal. Once again this is due to compensatory increases in Complex I, III, and IV activity, with their fluxes doubled in the worst case. Unfortunately we also observe the corresponding effects, as noted in Section 3.2, on oxygen consumption. As noted before, oxygen consumption which is reported in uncoupling in Table 6 is directly proportional to Complex IV activity.

### 3.8 Oxidative Phosphorylation and Mitochondrial Disease in the Proximal Tubule

Mitochondrial diseases, as well as the consequences of certain drugs like NRTIs, do not have a single universal effect on oxidative phosphorylation. For instance, oxidative stress can cause any combination of OXPHOS dysfunctions, and that is a key pathway for NRTI-induced mitochondrial dysfunction. We considered combinations of reductions by quarters in the activity of Complex I, Complex III, Complex IV, and ATP Synthase, which are the components of oxidative phosphorylation encoded by the mtDNA (and thus most at risk). We examined the steady state electrical potential gradient and cytoplasmic ATP concentration because the former maintains hydrogen flux through the electron transport chain and the latter is predictive of downstream loss of cell viability [58]. When noteworthy, ATP generation and oxygen consumption are also reported. Importantly, the P/O ratio remains highly consistent under OXPHOS dysfunction alone, and so a decrease in ATP generation is typically accompanied by a nearly proportionate decrease in oxygen consumption. This is due to the coupling of Complex IV (which determines oxygen consumption) and ATP Synthase (which determines ATP generation). The P/O ratio is in some sense a measure of the stoichiometry of the two reactions combined, and so it is usually significantly impacted only when intermediate products are lost (protons crossing the inner membrane by other means than ATP Synthase, i.e. hydrogen leakage).

#### 3.8.1 Univariate Parameter Changes

Examining one parameter at a time yields insights of the relative importance of the activities of each of the four components considered. The full set of univariate results for ATP concentration in the cytoplasm can be found in Figure 9. We see Complex III is much more important than the rest, likely due to which step is rate-limiting. There’s almost no discernable consequences of varying the activity of ATP Synthase for example, whereas ATP concentrations are reduced by 24% and the electrical potential gradient is reduced by 5 mV when Complex III activity is reduced to one quarter of typical enzymatic activity. That there may be no direct impact of ATP Synthase on ATP concentrations is not altogether unexpected. Across all cases where cytosolic ATP is greater than 90% of baseline, the flux through ATP Synthase remains within 4% of typical. ATP Synthase activity doesn’t significantly impact the ATP Synthase flux because ATP Synthase is not limiting for mitochondrial respiration. This is illustrated by results for ATP Synthase inhibition in mice neurons [23]. In that experimental model while there are indirect effects of ATP Synthase inhibition on murine neuronal aerobic respiration, the likely mechanism is via increased reactive oxygen species production, which leads to the downregulation of aerobic respiration as a whole, unrelated to the reduction in activity of ATP Synthase. A similar mechanism likely underlies Complex I-mediated mitochondrial dysfunction, it is well-known that it is a site of enhanced reactive oxygen species production during Complex I inhibition and that Complex I inhibition must be very strong before cellular respiration is inhibited [6, 72]. Low sensitivity to Complex I inhibition has in fact been observed in proximal tubule cell lines specifically [73].

**Figure 9:**
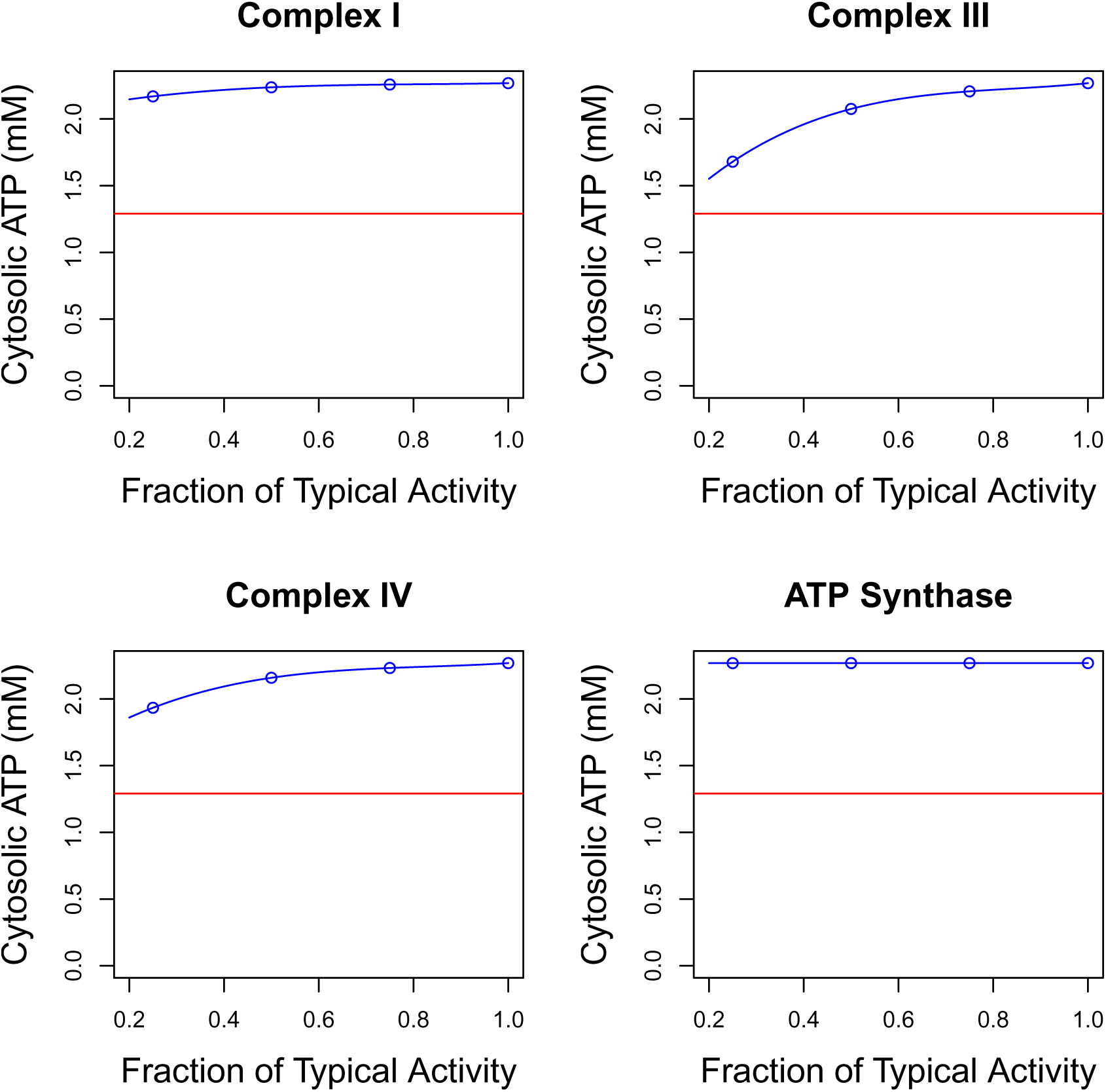
The effects on the cytoplasmic ATP concentration of changes to the activity of the four oxidative phosphorylation components considered in the PT. The red line represents the lowest ATP concentration found in any of the mitochondrial disease simulations performed.

Respiration has been shown to have similarly weak sensitivity to Complex IV activity in the rat kidney [55]. Theoreticians have long believed that the majority of respiratory enzymes are abundant far beyond what is necessary to maintain oxidative phosphorylation activity [55], and our predictions for ATP Synthase, Complex I, and Complex IV are compatible with that. Complex III inhibition was the largest exception in our model as noted above. Cytosolic ATP concentrations were predicted to be the most sensitive to Complex III inhibition by our model. The numbers for Complex III do not diverge too strongly from the literature however, in a study on a cell line of myoblasts, Schirris et al. observed that a roughly 75% reduction in Complex III activity produced a roughly 20% reduction in ATP generation [70]. In our model on the other hand there is a 10% reduction in ATP generation, which given the fact that we are working in a different tissue, and the heart has been observed to be more sensitive to inhibition of other OXPHOS enzymes [55] is not surprising.

#### 3.8.2 Multivariate Parameter Changes

We consider multivariate changes to oxidative phosphorylation due to the broad effects of Mitochondrial diseases on oxidative phosphorylation, as discussed later in Section 4. In considering these we wished to answer two questions: how much do multivariate changes extend the range of possible outcomes, and are there non-additive effects of interactions on the outcomes of interest? The first can be answered by comparing the range of outcomes from the univariate case to the multivariate case, using Figures 10. We see that Complex III still produces the most significant effects, explaining more of the variation in ATP concentration, as would be expected from the univariate case. What we see as well is that the added effect of combining activity reductions doesn’t significantly impact the electrical potential gradient, although the ATP concentration seems more sensitive.

**Figure 10:**
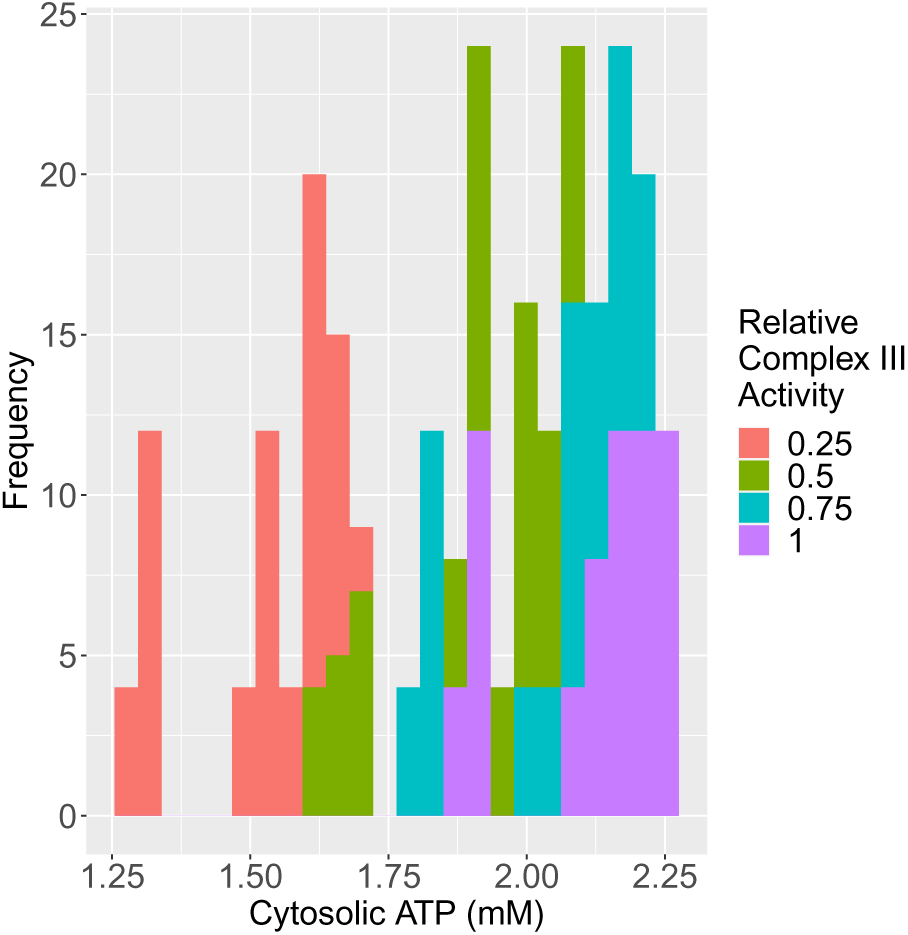
The cytosolic ATP concentration in each of the 256 cases considered, coloured by the activity of Complex III, after considering all the activities in the same manner as this graph we find that Complex III remains by far the most important determinant of the cytosolic ATP concentration.

To determine if there are non-additive interactions between the activities that were varied, a linear regression was used. Using least squares fitting, an affine plane was chosen as a function either of the univariate activities (our first model) or of the univariate activities combined with all the multiplicative interactions between them (our second model), in order to predict the cytosolic ATP concentration and the electrical potential gradient. The non-additive model performed no better than the univariate linear model. The *R*^2^ was almost the same for the two models, or in other words they fit the results almost equally well. The results of this analysis are found in Table 7.

**Table 7:** Models of cytosolic ATP concentrations and electrical potential gradients in the PT and mTAL, and the models’ R^2^ value. In the basic model, the only effects were additive in the relative change in the activity of each of the included OXPHOS components, in the interaction model, all combinations of multiplicative two-, three-, and four-way interactions were considered, which produced no meaningful difference in the model’s explanatory power.

| Tubule | Predicted Quantity | $R^2$ of Basic Model | $R^2$ of Model with All Interactions |
| --- | --- | --- | --- |
| PT | $[ATP_c]$ | 0.88 | 0.88 |
| PT | $\Delta\Psi$ | 0.91 | 0.91 |
| mTAL | $[ATP_c]$ | 0.84 | 0.84 |
| mTAL | $\Delta\Psi$ | 0.90 | 0.92 |

#### 3.8.3 Ifosfamide and Dichloroacetate

Ifosfamide reduces Complex I activity by half, and dichloroacetate has the capacity to prevent allosteric inhibition of pyruvate dehydrogenase in a manner that might compensate for respiratory dysfunction [28, 29]. The results for these cases, with the strongest considered case for the effect of dichloroacetate shown (a doubling of *K_i_*_NADH_), are in Figure 11 for cytosolic ATP concentration. Ifosfamide is nephrotoxic, and is specifically known to cause Fanconi syndrome. What the results suggest, compatible with our discussion above of Complex I existing in excess [6], is that the mechanism of nephrotoxicity may involve more than a direct loss of respiratory capacity. For instance, when Complex I is dysfunctional it may produce more reactive oxygen species, causing oxidative stress. Dichloroacetate shows little useful effect in these results, a result that matches the outcomes from experimental studies on dichloroacetate use with ifosfamide [62].

**Figure 11:**
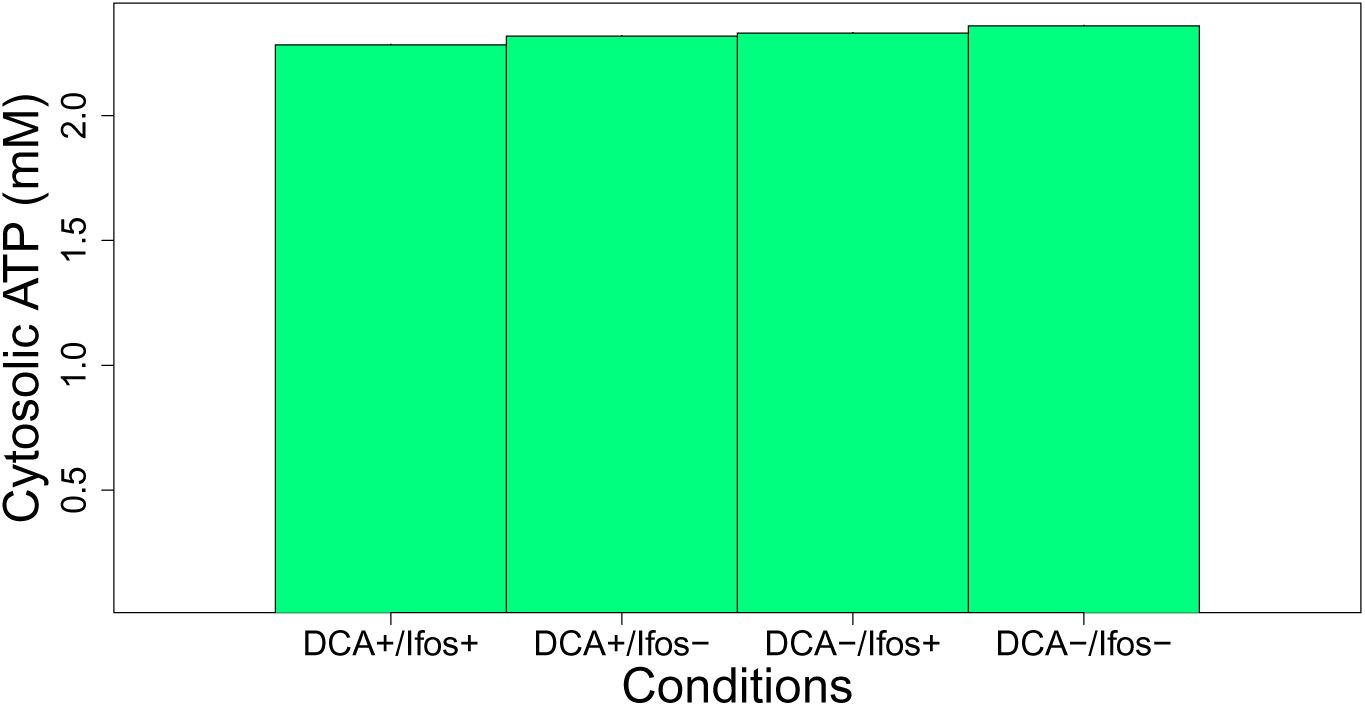
The effects of ifosfamide and dichloroacetate on cytosolic ATP concentration (mM) are shown.

#### 3.8.4 More General Drug Simulations of Nephrotoxic Drugs in the PT Mitochondria

Many drugs impact the general regulation of mitochondria in the cell and cause mitochondrial fragmentation. This tends to produce a generalized OXPHOS dysfunction and uncoupling. We represent this through combinations of uncoupling and OXPHOS dysfunction effects. This represents a more general set of combinations of the effects observed in mitochondrial disease, and as noted in Table 4. For instance Gentamicin, which acts as an uncoupler and generates reactive oxygen species that subsequently reduce OXPHOS activity [63, 65, 84].

Considering this set of cases essentially means adding another dimension of uncoupling (increased hydrogen leak activity) to the already considered OXPHOS parameter variations. The results for all such cases, with a maximum increased activity of hydrogen leak of 10 times normal, are shown in Figure 12 for the cytosolic ATP concentration. Another factor of interest to us is oxygen consumption, however while uncoupling appeared to increase oxygen consumption, OXPHOS dysfunction typically decreased oxygen consumption and so the effects never exceeded the effect on oxygen consumption of uncoupling alone.

**Figure 12:**
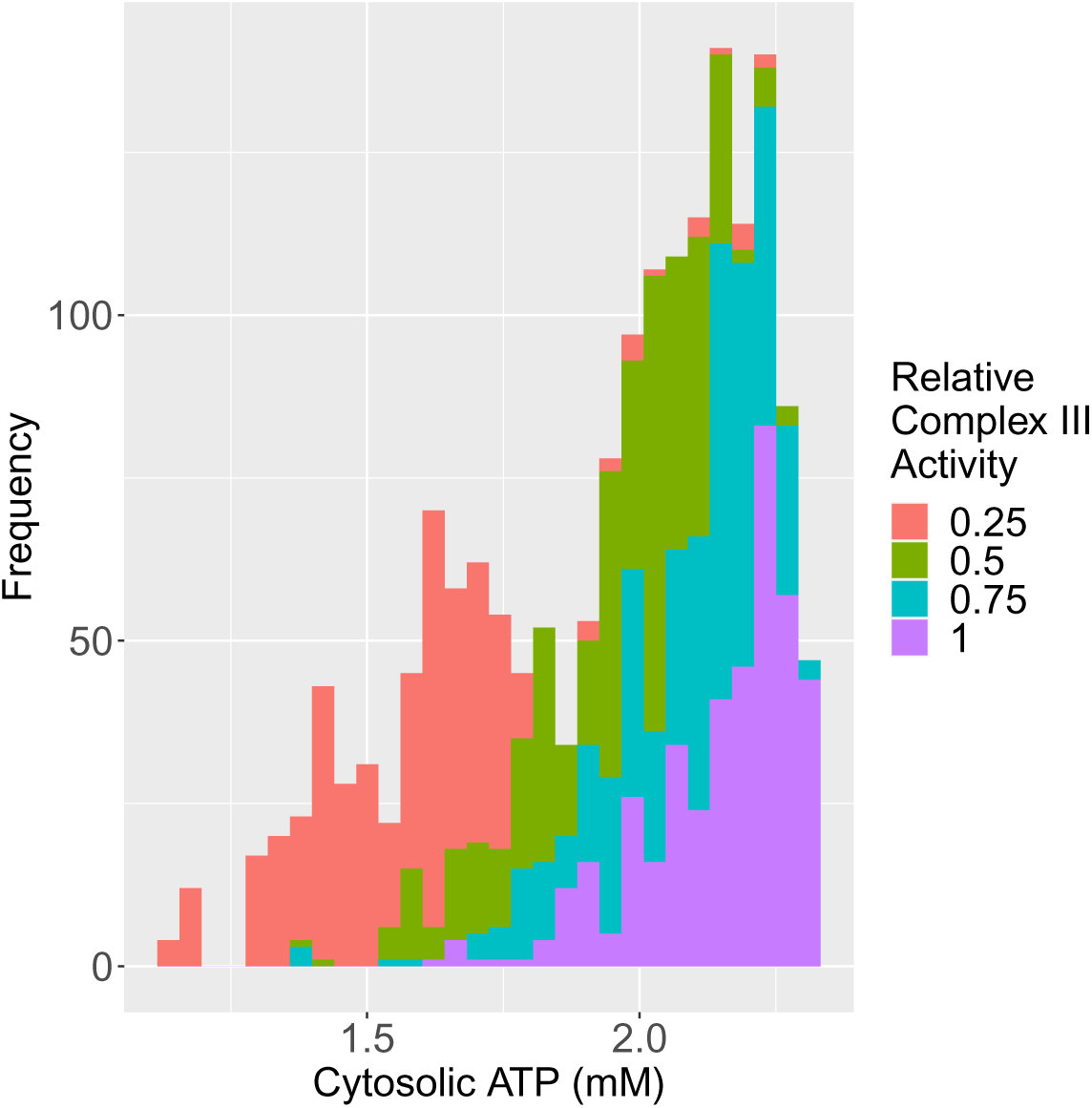
The ATP concentration in each of the cases considered of both OXPHOS dys-function and increased uncoupling, coloured by the relative activity of Complex III, after considering all of the activities we find that Complex III remains the most important determinant of the ATP concentration.

The results seen here show that the effect of Complex III is still extremely significant, but unlike in some previous cases, the lowest-activity case for Complex III is no longer disjoint from the other simulated cases. We now see that in some cases the combined effects on other activities can have as large of an effect on the ATP concentration and electrical potential gradient. This is evident from a linear model fit which predicts that the combined effects of the other variables will together have a comparable effect to that of Complex III activity reductions on their own.

### 3.9 Oxidative Phosphorylation and Mitochondrial Disease in the medullary Thick Ascending Limb

Here we followed the same procedure as in the PT and considered cases of combinations of OXPHOS activity reductions in the mTAL.

#### 3.9.1 Univariate Parameter Changes

The univariate cases for ATP concentration show large changes in the cytosolic ATP concentration under OXPHOS dysfunction, relatively comparable to the sensitivity predicted above in the PT. The mTAL appears to be most sensitive to dysfunction of Complex IV, unlike the PT which is most sensitive to Complex III dysfunction. The mTAL is also sensitive to Complex III dysfunction like the PT. We see that the mTAL is more sensitive to Complex IV dysfunction than the PT is. This different prediction can be explained simply: the activity of Complex IV is limited by the far lower oxygen tension in the mTAL. When we consider comparable Complex IV dysfunction (75% lower Complex IV activity) in the mTAL, but now with a PT-like 50 mmHg oxygen tension, we do not see the same sensitivity to Complex IV dysfunction (no change in cytosolic ATP concentrations in the 50 mmHg oxygen tension case, compared to a 31% change in cytosolic ATP concentrations in our baseline 10 mmHg oxygen tension case). Otherwise the results strongly resemble the PT case.

#### 3.9.2 Multivariate Parameter Changes

Like for the PT we wish to address whether there are non-additive effects, and the total range of possible outcomes under multiple OXPHOS complex deficiencies for the mTAL, thereby capturing many mitochondrial diseases which often include these combined effects. In Figure 15 we show the range of cytosolic ATP concentrations for various combinations of OXPHOS dysfunction. Once again we fit a linear model and a model with multiplicative interaction terms, and like for the PT, the *R*^2^ is above 0.8 for the linear models and the improvement from including interactions is marginal. The results can once again be found in Table 7.

**Figure 13:**
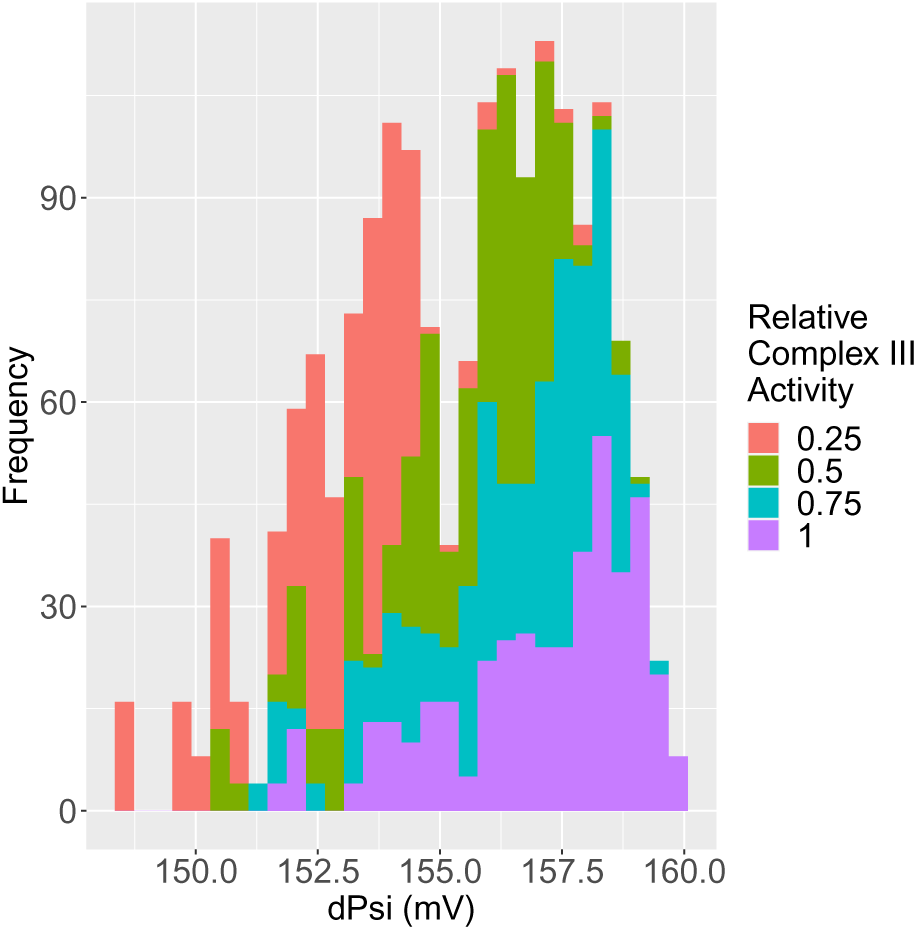
The electrical potential gradient in each of the cases considered of both oxidative phosphorylation dysfunction and increased uncoupling, coloured by the activity of Complex III, after considering all of the activities in the same manner as this graph we find that Complex III remains the most important determinant of the electrical potential gradient.

**Figure 14:**
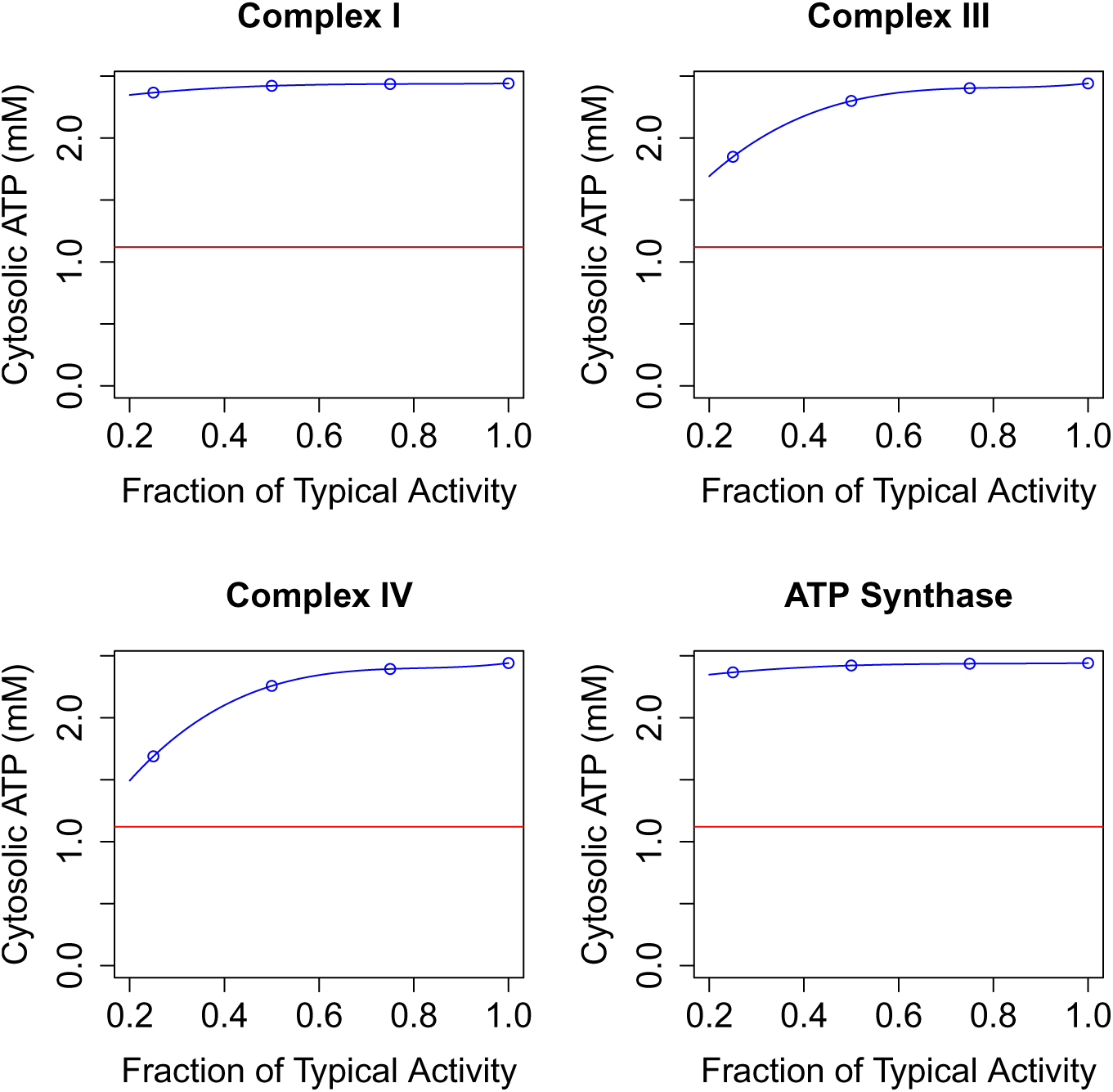
The effects on the cytoplasmic ATP concentration of changes to the activity of the four oxidative phosphorylation components considered in the mTAL. The red line represents the lowest ATP concentration found in any of the mitochondrial disease simulations performed.

**Figure 15:**
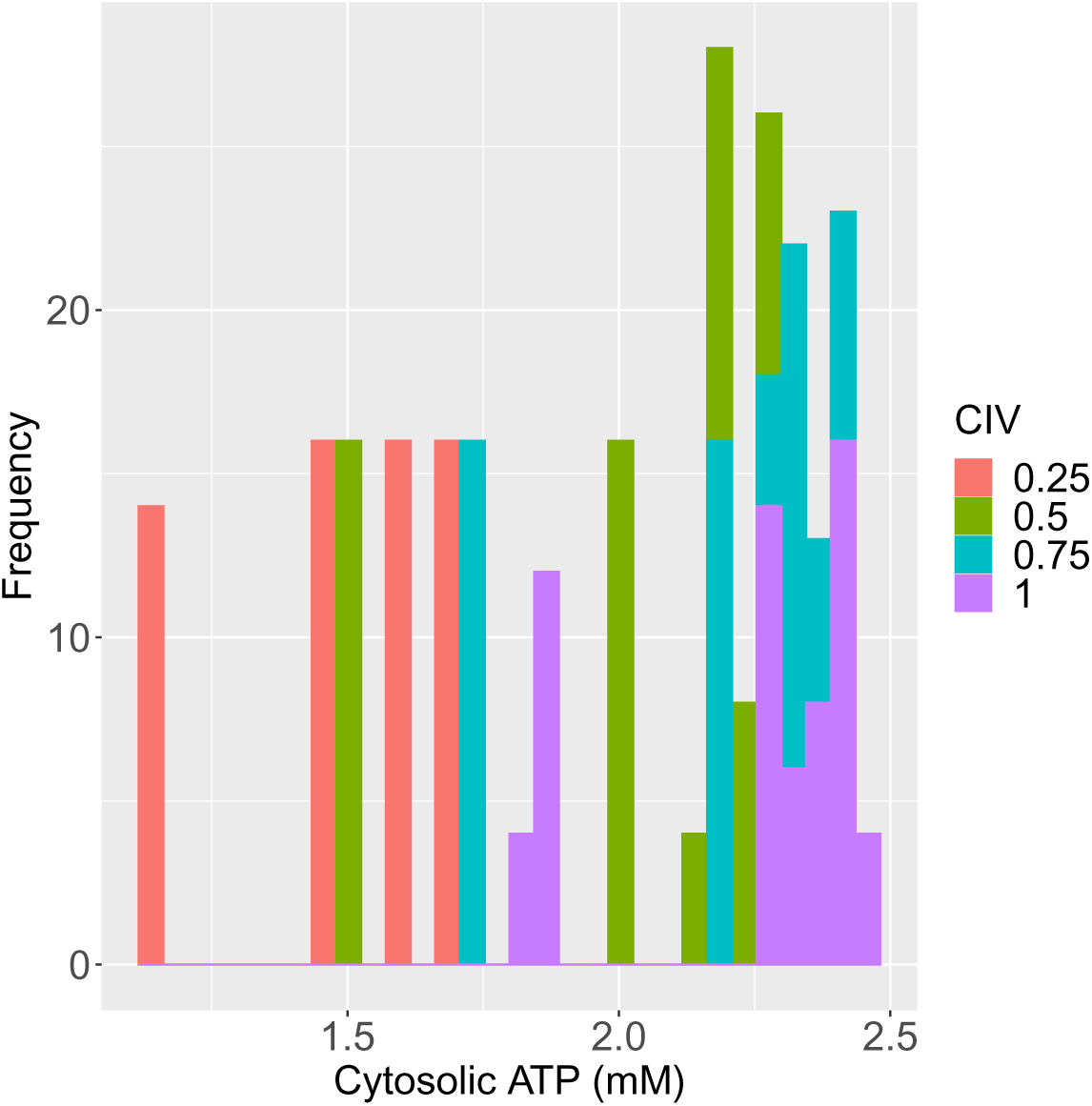
Cytosolic ATP levels in the mTAL in our mitochondrial disease simulations (various combinations of OXPHOS dysfunction), coloured by the level of Complex IV activity relative to typical Complex IV activity.

#### 3.9.3 OXPHOS Deficiencies Combined with Other Effects

OXPHOS dysfunction as previously discussed, is often concurrent with uncoupling, such as in diabetes, and we know the mTAL to be especially prone to hypoxic injury due to low tissue oxygen tension. Thus we considered these two factors as well to see their effect in combination with OXPHOS deficiency. We find that significant drops in oxygen tension lead to the largest effect in these combined cases. Under these combined effects, as shown in Figure 16, we still predict that Complex IV activity is a particularly significant parameter for the maintenance of cytosolic ATP levels.

**Figure 16:**
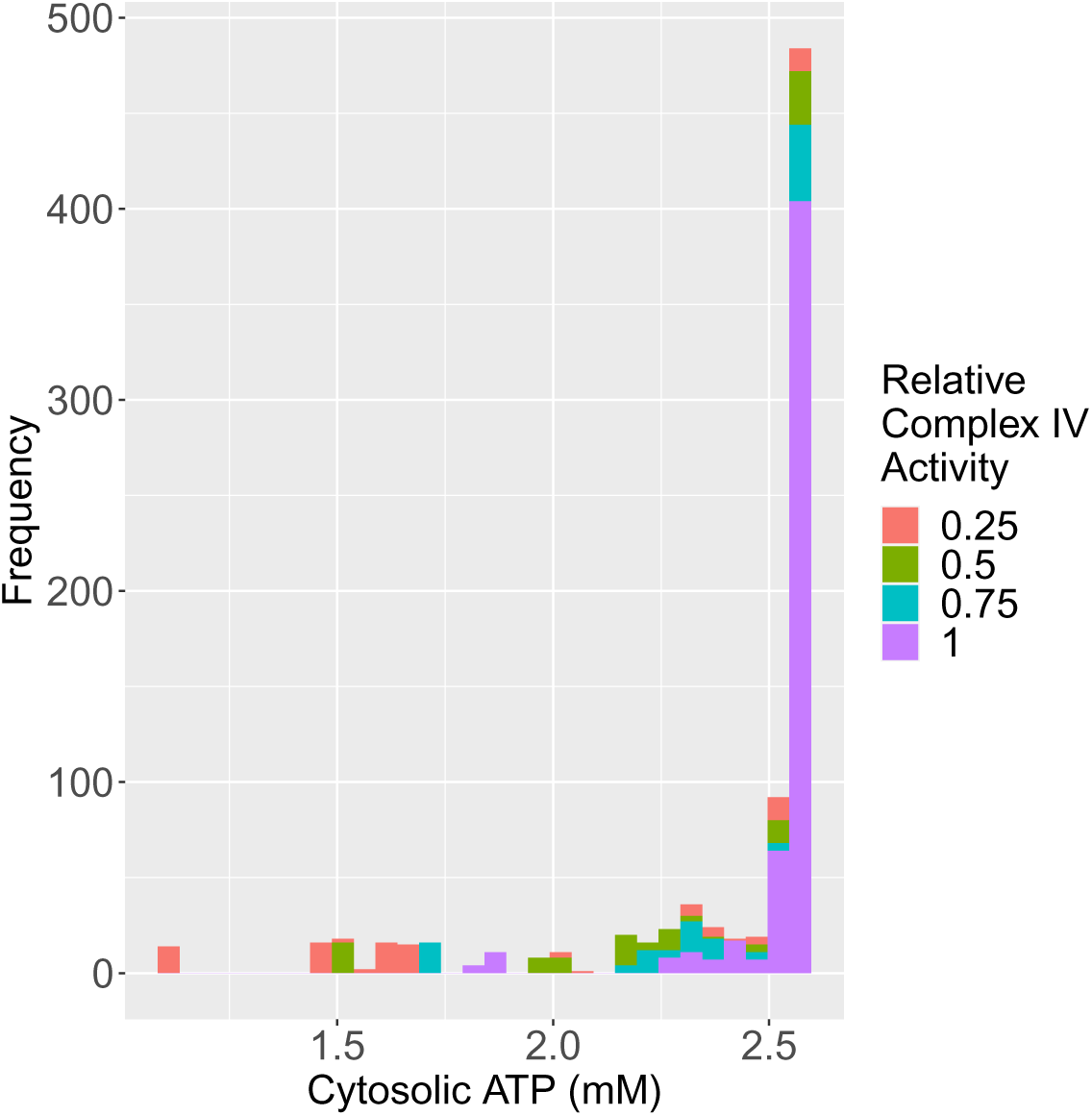
The cytosolic ATP levels for varied OXPHOS dysfunction, uncoupling, and oxygenation, coloured by Complex IV activity levels.

### 3.10 Hypoxia in Proximal Tubule Mitochondria

The kidney is sensitive to reductions in tissue oxygen partial pressure, and understanding the associated threshold for injury is both medically and physiologically significant. Micro- and macrovasculature problems may produce local hypoxia, often both are involved. The effects of macrovascular hypoxia though are more easily measurable, and we can rule out their sufficiency for hypoxic injury by determining whether the macroscale oxygen partial pressure is low enough to cause hypoxic injury, if it isn’t then microvasculature problems must be contributing on a smaller scale than is measured [11, 59]. The results of a gradation of oxygen partial pressure reductions are shown in Figure 17. From this we see a degree of insensitivity of cytosolic ATP concentrations to oxygen partial pressure dropping in the renal cortex until 5-10 mmHg, where we see a much greater drop in ATP concentrations.

**Figure 17:**
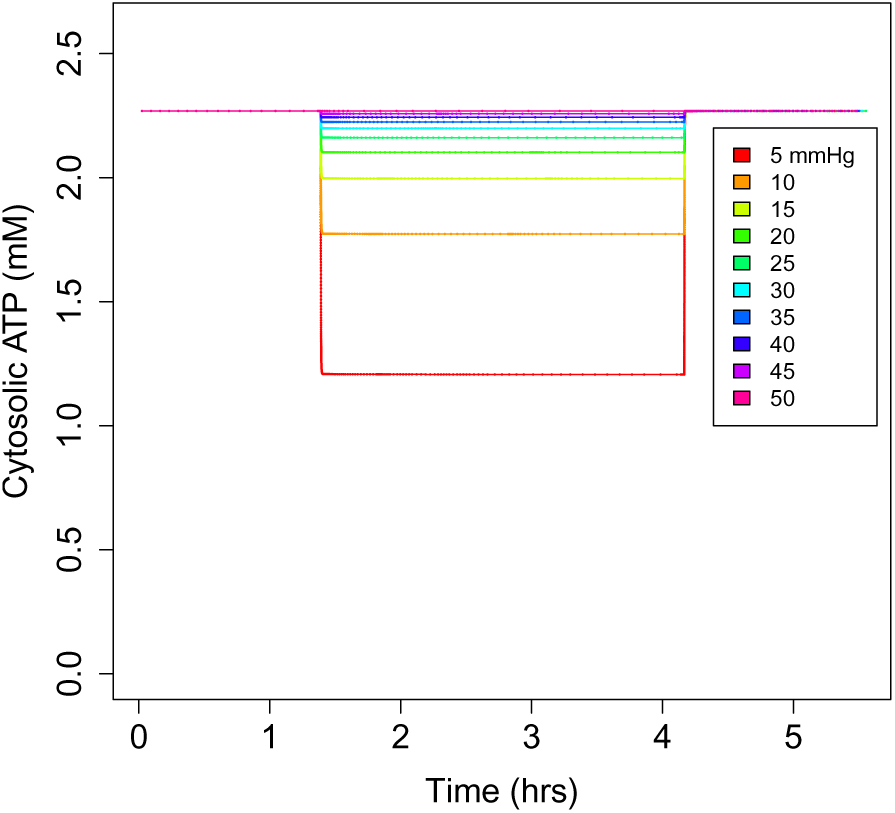
Cytosolic ATP concentration in the PT for a range of reductions by tenths of oxygen partial pressure (starting at 5000 s), to a lowest of one tenth of baseline oxygen partial pressure (or 5 mmHg) from a maximum of 50 mmHg. This is followed by reoxygenation at time 15,000 s.

Reoxygenation is also considered in the simulation shown in Figure 17, in our model, the predicted response is immediate and returns to the previous equilibrium.

The strong robustness to hypoxia, down to 10 mmHg in the proximal tubule (with significant changes in cell behaviour between 5 and 10 mmHg), agrees with measurements of a ‘critical’ *P_O_*_2_ found in experimental work of 10 mmHg [19]. This robustness is due to a sustained capacity to consume the necessary amount of oxygen until that point. For instance, at normoxic oxygen tensions (50 mmHg) we see an oxygen consumption of 2.47 nmol *O*_2_ per milligram per second in our model, and at 10 mmHg, the cell still consumes 2.30 nmol *O*_2_ per milligram per second in our model.

### 3.11 Hypoxia in medullary Thick Ascending Limb Mitochondria

A much smaller absolute change is required in order to produce an extremely low oxygen tension in the mTAL, because the baseline tissue oxygen tension of the mTAL is much lower than that of the PT (10-20 mmHg in the mTAL compared to 50 mmHg for the PT). However as noted by Schiffer, Gustafsson, and Palm [69], there are multiple adaptations that mTAL mitochondria have to address this problem, most notably: they are more densely packed in the mTAL, they have greater efficiency, and they have greater oxygen affinity [69]. These features are included in our model, and produce a partial robustness. The cytosolic ATP levels are greater than 2 mM until below 4 mmHg, whereas at 5 mmHg the PT’s cytosolic ATP levels much worse than necessary to kill the PT cell. Thus, the mTAL can survive much absolutely smaller oxygen tensions. However since the baseline oxygen tension is lower, it is not robust to comparably extreme variations of the oxygen tension, the PT’s oxygen tension may be as low as 20% of typical before being in major danger. The mTAL’s cytosolic ATP levels are much lower than baseline at oxygen tensions below 40% of typical. In Figure 18 we see the effects of adjusting the oxygen tension in the mTAL.

**Figure 18:**
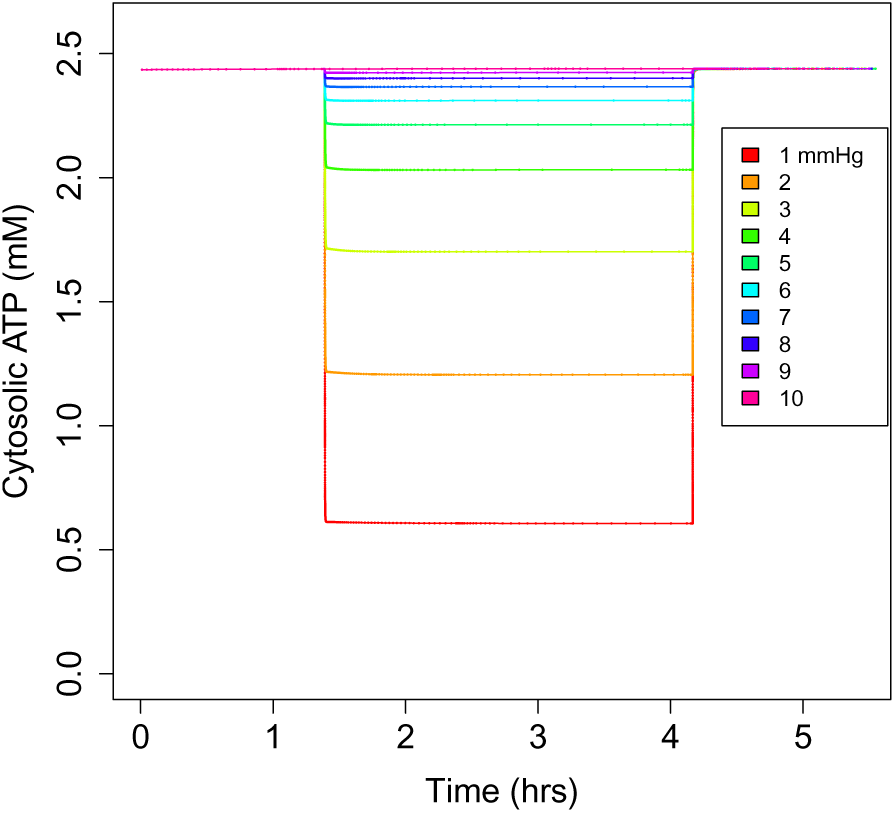
Cytosolic ATP concentration in the mTAL with high ATP consumption for a range of reductions by tenths of oxygen partial pressure (starting at 5,000 s), to a lowest of one tenth of baseline oxygen partial pressure (or 1 mmHg) from a maximum of 10 mmHg. This is followed by reoxygenation at time 15,000 s.

Like with OXPHOS dysfunction, we wished to identify the factors contributing to the robustness of the mTAL under hypoxia. To do this we consider 1 mmHg normal oxygen tension in the mTAL (1 mmHg) but with PT-like mitochondrial volume fraction or Complex IV oxygen affinity (and combinations thereof). We compare these cases with the PT’s response to 1 mmHg oxygen tension. We find that while a PT-like mitochondrial volume fraction makes the mTAL cell perform worse as expected, it is necessary and sufficient that you change the Complex IV oxygen affinity to PT-like values in order to produce comparable drops in the cytosolic ATP content in the mTAL. This indicates the pivotal role of Complex IV oxygen affinity in driving the mTAL’s robustness against hypoxia.

### 3.12 Ischemia-Reperfusion Injury in the Proximal Tubule

During ischemia, the cell suffers a severe decline in oxygen concentration. As a consequence of ischemia, the adenine nucleotide (AMP/ADP/ATP) pool becomes smaller, an effect we include. For our model of reperfusion, the oxygen concentration returns to normal and the combined adenine nucleotide pool is 30% of its typical size, based on the results of Cunningham, Keaveny, and Fitzgerald [14]. In the post-reperfusion steady state with this model, we predict the proton motive force is much high at 182 mV (compared to 165 mV typically). The NADH/NAD^+^, coenzyme Q and cytochrome C pools are not predicted to be left in a highly reduced state. Until the adenine nucleotide pool recovers the cell may be susceptible to reperfusion injury on these grounds. Adding dichloroacetate, which increases the flux through pyruvate dehydrogenase, did not significantly impact the redox state of the cell following reperfusion. Tripling the cytosolic pyruvate concentration, which also increases the flux through pyruvate dehydrogenase, similarly had no effect.

### 3.13 Ischemia-Reperfusion Injury in the medullary Thick Ascending Limb

Ischemia is represented in the same way for the mTAL as for the PT. Like for the PT, we have a highly reduced NADH/NAD^+^ pool and an elevated proton motive force at steady state. 80% of the NADH/NAD^+^ pool is NADH and the proton motive force is 186 mV. The coenzyme Q and cytochrome C pools are not left in a highly reduced state. Dichloroacetate and cytosolic pyruvate had small effects on the redox state of electron carriers and proton motive force.

## 4 Discussion

The goal of the present study was to develop a new model of mitochondria in the mTAL, to improve a model of mitochondria in the PT, and to examine both in several pathological conditions. Our model is able to predict several important cellular quantities, including ATP generation, P/O (Phosphate/Oxygen) ratio, proton motive force, electrical potential gradient, oxygen consumption, the redox state of important electron carriers, and ATP consumption. As noted above, for many of these quantities our predictions were within the range of known measurements both for *in vivo* and *in vitro* cases.

Schiffer et al. notes a lower P/O ratio in the PT than the mTAL, indicating that the mTAL is more metabolically efficient [69]. The P/O ratio is the ratio of ATP synthase phosphorylation of ATP to the Complex IV oxygen consumption. The ratio *in vitro* has been measured to be 1.93±0.13 in the mTAL, and 1.81±0.10 in the PT [69]. Our model predicts a similar *in vitro* P/O ratios between the PT and mTAL, but lower in the PT (see Table 2), like we see in Schiffer et al. The predicted P/O ratios *in vivo* are found in Tables 5 and 6 respectively, and we see that *in vivo* the mTAL has a marginally higher P/O ratio. This is driven *in vivo* by higher metabolic demand and lower oxygen consumption due to the mTAL’s comparatively hypoxic state.

Before we get into the details of the dysfunctional cases, we define certain well-understood adverse outcomes. ATP depletion is a significant drop in the cytosolic ATP concentration. ATP depletion is one pathway by which cells die. In mice, there is significant risk of apoptosis from ATP depletion below 70% of typical cytosolic ATP concentrations, and necrosis is assured below 15% [58]. These benchmarks help us to understand the severity of the ATP depletion observed under the conditions we simulate. Depolarization is a significant loss of either electrical potential gradient, pH difference, or both across the inner membrane of the mitochondria. Remaining polarized is a necessary prerequisite for ATP generation by the mitochondria in a way that is included in our model or the model equations surrounding the electron transport chain). At equilibrium, ATP generation matches consumption. Reduced ATP generation will limit how much ATP may be consumed, leading to poor reabsorption on the part of the kidney. Finally, overproduction of reactive oxygen species is understood to be a major problem for instance in ischemia-reperfusion injury. We capture this in our model by examining the reduction state of the NADH, Cytochrome C, and co-enzyme Q pools in the cell. For instance, if there is much more NADH than NAD^+^, then Complex I will be in a more reduced state, and if simultaneously there is more of the reduced form of coenzyme Q than its oxidized form, then Complex I may create reactive oxygen species. Similarly Complex III may do the same if the pools of co-enzyme Q and Cytochrome C are both in a reduced state.

Model simulations indicate that the PT and mTAL are sensitive to different metabolic perturbations, we deal with these pathological questions first. Both are predicted to be susceptible to a lowered activity of Complex III, but the mTAL is more susceptible to Complex IV deficiency. This result is unsurprising because Complex IV uses oxygen as a substrate, and the relatively hypoxic environment of the mTAL means that a larger amount of Complex IV is necessary in order to produce the same hydrogen flux. On the other hand though, if we consider the two tissues’ sensitivities to hypoxia in absolute terms we see that at an oxygen tension of 5 mmHg, the cytosolic ATP concentration in the mTAL is comparable to the baseline cytosolic ATP concentration in the PT. When the PT experiences an oxygen tension of 5 mmHg, it experiences ATP depletion sufficient to cause cell death. This is possible because Complex IV has a higher oxygen affinity in the mTAL in our model and experimentally [69]. That said, if we consider hypoxia relative to baseline oxygen tensions in each tissue, we see that the mTAL is at risk of cell death for 70% declines in the baseline oxygen tension whereas the PT is not at risk of cell death until 80-90% declines in oxygen tension from baseline. If we consider instead ATP homeostasis for higher rates of ATP consumption, we see that for a *Q_ATP,max_* that is 30% higher than typical in the mTAL (see the highest *Q_ATP,max_* in Table 6), we have a higher cytosolic ATP concentration than the PT does even after only a 25% increase in the maximal ATP consumption (see Table 5). This is despite the fact that the baseline ATP consumption in the PT is much lower than in the mTAL. In fact, after a 25% increase in the maximal ATP consumption in the PT, the PT’s maximal ATP consumption is still lower than the baseline maximal ATP consumption in the mTAL. Ultimately then the PT is more sensitive to hypoxia below 10 mmHg oxygen tension and to increased metabolic demand. The mTAL is more sensitive to equivalent proportional decreases in oxygen tension than the PT is, and to Complex IV dysfunction. The mTAL is more sensitive to hypoxia and Complex IV dysfunction because of its baseline low oxygen tension, which can be demonstrated by considering its cytosolic ATP concentrations at comparable oxygen tensions to our PT hypoxia cases. The PT is more sensitive to oxygen tensions below 10 mmHg because Complex IV oxygen affinity in the mTAL is greater.

The proximal tubule is susceptible to OXPHOS dysfunction, for instance due to the accumulation of drugs reabsorbed from the nephron lumen [31]. Our results suggest that in the proximal tubule of the rat kidney, ATP concentrations are most sensitive to Complex III activity reductions, as can be seen in Figure 9. Based off of our results, we can predict that recovering Complex III function is the most promising target for mitochondria-mediated Fanconi syndrome if Complex III is affected. The predicted importance of Complex III, is compatible with experimental studies that suggest its importance to mitochondria-mediated renal pathology [61]. The additive relationship noted above between distinct OXPHOS deficiencies is indicative that this advice will remain true regardless of the combination of OXPHOS deficiencies.

Our model doesn’t directly predict the production of reactive oxygen species, as we’ve noted before. However what it does predict presents an interesting case for how out-of-control reactive oxygen species production may depend on the supply of adenine nucleotides available during reperfusion. When ADP is not available within the mitochondrion, ATP Synthase is not able to shuttle hydrogen ions across the inner membrane of the mitochondrion. Without this, the electron transport chain is left in a reduced state and the proton motive force is increased. The adenine nucleotide levels (that is, the pooled concentration of ATP, ADP, and AMP) are experimentally observed to fall to roughly 30% of typical after clamping the renal pedicle (which includes the renal artery) for 60 minutes [14]. With a lower adenine nucleotide level, we see that the reperfused kidney’s mitochondria are left in a reduced state in the PT and mTAL in our model as we expected, as discussed in Sections 3.12 and 3.13. This highly reduced state persists in our model unless the adenine nucleotide levels recover. The size of the adenine nucleotide pool is well-understood to be important to the respiration rate because adenine nucleotide translocase activity is frequently rate-limiting for mitochondrial respiration [4] and the size of the adenine nucleotide pool controls the flux through adenine nucleotide translocase [22]. Since it is important to the respiration rate, it is natural that with a smaller adenine nucleotide pool, that earlier steps in oxidative phosphorylation would be left in a reduced state for lack of a target to donate their electrons to.

Previous work [19] and our model predictions show that under 10 mmHg PT mitochondrial matrix oxygen tension, and our work predicts under 4 mmHg mTAL mitochondrial matrix oxygen tension, that some cells will experience ATP depletion. These estimates are much lower than tissue oxygen tensions linked to hypoxia [26], indicating that the hypoxia is more local than tissue-based measures can find, a result highly suggestive of microvascular ischemia, in accordance with the conventional microvascular view of renal hypoxia [21, 53, 59]. This suggests a role for microvascular ischemia in the development of renal hypoxia, rather than for macrovascular effects on renal hypoxia alone. Our findings for the threshold of hypoxic injury in the proximal tubule are compatible with experimental results [19].

We note above that for a physiologically reasonable metabolic demand, the mTAL can be sensitive to hypoxia. Understood relative to the PT’s oxygen tension though, the mTAL is fairly resilient. We wish to identify the source of the mTAL’s resilience. We identified three factors in Section 3.11 which contribute to the resilience of the mTAL to ATP depletion during hypoxia: the mitochondrial volume fraction and metabolic demand like in the previous case of resilience that we were interested in, and Complex IV’s oxygen affinity, which along with the above parameters is recognized by physiologists as important to the resilience of the mTAL [69]. Our results noted in Section 3.11 tell a nuanced story as to what provides the mTAL its robustness. For a 90% decrease in oxygen tension in the PT, we saw an 81% decrease in cytosolic ATP concentration. If we only change one of the above identified parameters, the worst change in cytosolic ATP concentrations we may produce is a 27% change from increasing the metabolic demand. However by combining PT-like metabolic demand with either PT-like mitochondrial volume fraction or Complex IV oxygen affinity, we can make the mTAL more sensitive to hypoxia than the PT. Thus the robustness of the mTAL can be explained by all three factors noted above, once again the metabolic demand is important, but altering either Complex IV oxygen affinity or the mitochondrial volume fraction as well to PT-like values is enough to remove the mTAL’s robustness against hypoxia. While experimental work has started to attend to Complex IV’s oxygen affinity [69], this work is incomplete and we hope for direct estimates of Complex IV’s oxygen affinity in the future. Our work uses Schiffer et al’s [69] measurements of mitochondrial oxygen consumption to fit the oxygen affinity of Complex IV, but more direct estimates would increase our confidence in our predictions. Our model predictions corroborate Schiffer et al’s [69] hypothesis about the basis for the mTAL’s resilience against hypoxia: that the mitochondrial volume density and Complex IV oxygen affinity both contribute to the robustness of the mTAL. This goes towards answering our comparative questions about the PT and mTAL.

Our predictions in our drug simulations indicate that the circumstances under which an uncoupler such as salicylate could cause proximal tubulopathy in rats are limited. We found that uncoupling was not a significant cause of ATP depletion in the cell in our model, although it could cause a significant increase in oxygen consumption (see Sections 3.6 and 3.7 for the results of the uncoupling case). Likely it requires underlying pre-existing conditions in the rat, for instance affecting delivery of oxygen to the kidney, such that the uncoupler is a more potent cause of hypoxia. Similarly, ifosfamide appears to be a less potent cause of kidney injury in rats according to our model (see Section 3.8.3). Ifosfamide primarily affects Complex I, and at experimentally noted ifosfamide-caused reductions in Complex I activity we did not observe significant ATP depletion in our model. This accords with observations that ifosfamide’s nephrotoxicity is more concerning in children and patients taking cisplatin concurrently [40]. Ifosfamide could also produce an indirect impact through oxidative stress, a known consequence of Complex I dysfunction and ifosfamide use that is not within the scope of our model [72]. On the other hand, gentamicin is known to have common nephrotoxic effects via several mitochondrial mechanisms, including OXPHOS dysfunction and uncoupling [66], and our model indicates that these are sufficient to cause massive ATP depletion (see Section 3.8.4 and 3.9.3).

Aside from the above potential adverse outcomes, we studied oxygen consumption and how it was affected by many factors but principally uncoupling. Particularly in diabetes, uncoupling reduces the efficiency of aerobic respiration and increases the kidney’s oxygen consumption [26]. Our work predicts increased oxygen consumption, but cannot predict the subsequent hypoxia because it treats mitochondrial partial pressure of oxygen as a clamped quantity, and thus not responsive to oxygen consumption. Future experimental and modelling work could go further by tying oxygen consumption to the partial pressure of oxygen via an estimated response curve of oxygen tension to oxygen consumption or a control ratio of the two quantities. However with this assumption of a clamped oxygen tension, we noted that a 10x increase in hydrogen leak produced a 39% increase in oxygen consumption in the PT, and that a 10x increase in hydrogen leak in produced a 28% increase in oxygen consumption in the mTAL. In light of the mTAL’s inelastic supply of oxygen [19] and low baseline oxygen tension, it may nevertheless be more susceptible to uncoupling-mediated hypoxia.

In conclusion, our model accurately captures mitochondrial ATP generation and oxygen consumption in the PT and mTAL. This model allows us to measure the efficiency of aerobic respiration in these two tissues, and the capacity of mitochondria to meet metabolic demand in the PT and mTAL. These predictions, found in Table 5 and 6, match well with experimental observations. We’ve also simulated OXPHOS dysfunction, uncoupling, hypoxia, and ischemia-reperfusion. Where possible we’ve identified thresholds for ATP depletion based on the above kinds of dysfunction. In the case of ischemia-reperfusion, we identify how to indirectly examine the production of reactive oxygen species from our model. We also examine different model descriptions of the process of reperfusion, demonstrating that the risk of reactive oxygen species production is much greater under some descriptions of reperfusion than others. More accurately capturing the feedback between tissue oxygen tension and oxygen consumption would be a valuable next step in capturing the negative consequences of uncoupling and other mitochondrial dysfunctions.

Ultimately then, our results indicate that each of the tissues we considered have characteristic ways that they are sensitive to mitochondrial dysfunction, or mitochondria-mediated injury. These ways can be connected to a range of mitochondrial and tissue-scale factors. For instance, the activity of OXPHOS enzymes is crucial to the robustness of the liver and is a mitochondrial-scale factor. On the other hand, tissue-scale factors such as high metabolic demand (necessary in the kidney for instance for urine concentration [5]) and oxygen tension appear in our model to be crucial for understanding the risks from mitochondrial diseases (once again see Section 3.9). This opens up space for extending our model: we know that the tissue may affect the mitochondria, but in aggregate, mitochondria may also affect the tissue. A limitation of our model is that tissue-scale factors are not modelled as dynamically responsive to mitochondrial functioning. For example, oxygen tension is clamped in our model, meaning that regardless of local oxygen consumption the tissue’s oxygen tension remains the same. Consequently, if we want to understand the effect of a certain degree of uncoupling (which increases oxygen consumption), we have no better method of pursuing this than simply seeing how it affects the mitochondria to combine that level of uncoupling with various degrees of hypoxia, without any way of predicting the appropriate degree of hypoxia. This limitation points towards a corresponding ambition however: multi-scale modelling of metabolism. We believe the best way to expand on this work is to integrate it into larger models of epithelial transport [37, 45, 38, 36, 48, 50, 47, 49, 46, 57, 18, 44] and oxygenation [75, 74, 12, 13, 27] and of the factors determining local metabolic demand. Appropriate multi-scale models of metabolism will involve more than just placing our mitochondrial models inside of a larger scale (and ultimately, hopefully whole-body) model with appropriate interfaces. While the liver and kidney are particularly important to whole-body homeostasis, every tissue has mitochondria and most tissues are susceptible to mitochondrial dysfunction or mitochondria-mediated injury. For a whole-body multiscale model of metabolism then, we need models of mitochondrial respiration in other tissues.

## References

[1] Alston Charlotte L, Rocha Mariana C, Lax Nichola Z, Turnbull Doug M, and Taylor Robert W. The genetics and pathology of mitochondrial disease. The Journal of pathology 241(2): 236–250, 2017.

[2] Angelini Corrado, Bello L, Spinazzi M, and Ferrati C. Mitochondrial disorders of the nuclear genome. Acta Myologica 28(1): 16, 2009.

[3] Assmann Nadine, Dettmer Katja, Simbuerger Johann MB, Broeker Carsten, Nuernberger Nadine, Renner Kathrin, Courtneidge Holly, Klootwijk Enriko D, Duerkop Axel, Hall Andrew M, and others. Renal fanconi syndrome is caused by a mistargeting-based mitochondriopathy. Cell Reports 15(7): 1423–1429, 2016.

[4] Atlante Anna, Seccia Teresa Maria, Marra Ersilia, and Passarella Salvatore. The rate of atp export in the extramitochondrial phase via the adenine nucleotide translocator changes in aging in mitochondria isolated from heart left ventricle of either normotensive or spontaneously hypertensive rats. Mechanisms of ageing and development 132(10): 488–495, 2011.

[5] Aw Mun, Armstrong Tamara M, Nawata C Michele, Bodine Sarah N, Oh Jeeeun J, Wei Guojun, Evans Kristen K, Shahidullah Mohammad, Rieg Timo, and Pannabecker Thomas L. Body mass-specific na+-k+-atpase activity in the medullary thick ascending limb: implications for species-dependent urine concentrating mechanisms. American Journal of Physiology-Regulatory, Integrative and Comparative Physiology 314(4): R563– R573, 2018.

[6] Barrientos Antoni and Moraes Carlos T. Titrating the effects of mitochondrial complex i impairment in the cell physiology. Journal of Biological Chemistry 274(23): 16188– 16197, 1999.

[7] Benard Giovanni, Faustin Benjamin, Passerieux Emilie, Galinier Anne, Rocher Christophe, Bellance Nadege, Delage J-P, Casteilla Louis, Letellier Thierry, and Rossignol Rodrigue. Physiological diversity of mitochondrial oxidative phosphorylation. American Journal of Physiology-Cell Physiology 291(6): C1172–C1182, 2006.

[8] Bouby Nadine, Bankir Lise, Trinh-Trang-Tan Marie-Marcelle, Minuth Will W, and Kriz Wilhelm. Selective adh-induced hypertrophy of the medullary thick ascending limb in brattleboro rats. Kidney international 28(3): 456–466, 1985.

[9] Brand Martin D and Nicholls David G. Assessing mitochondrial dysfunction in cells. Biochemical Journal 435(2): 297–312, 2011.

[10] Budunova Irene V and Mittelman Leonid A. The effect of k+/h+ antiporter nigericin on gap junction permeability. Cell biology and toxicology 8(1): 63–73, 1992.

[11] Calzavacca Paolo, Evans Roger G, Bailey Michael, Bellomo Rinaldo, and May Clive N. Cortical and medullary tissue perfusion and oxygenation in experimental septic acute kidney injury. Critical care medicine 43(10): e431–e439, 2015.

[12] Chen Jing, Edwards Aurélie, and Layton Anita T. Effects of ph and medullary blood flow on oxygen transport and sodium reabsorption in the rat outer medulla. American Journal of Physiology-Renal Physiology 298(6): F1369–F1383, 2010.

[13] Chen Jing, Sgouralis Ioannis, Moore Leon C, Layton Harold E, and Layton Anita T. A mathematical model of the myogenic response to systolic pressure in the afferent arteriole. American Journal of Physiology-Renal Physiology 300(3): F669–F681, 2011.

[14] Cunningham SK, Keaveny TV, and Fitzgerald P. Effect of allopurinol on tissue atp, adp and amp concentrations in renal ischaemia. Journal of British Surgery 61(7): 562–565, 1974.

[15] Dowd T, Barac-Nieto M, Gupta RK, and Spitzer A. 31p nuclear magnetic resonance and saturation transfer studies of the isolated perfused rat kidney. Kidney and Blood Pressure Research 12(3): 161–170, 1989.

[16] Dzbek Jaroslaw and Korzeniewski Bernard. Control over the contribution of the mitochondrial membrane potential (*δψ*) and proton gradient (*δ*ph) to the protonmotive force (*δ*p): in silico studies. Journal of Biological Chemistry 283(48): 33232–33239, 2008.

[17] Edwards A, Palm F, and Layton AT. A model of mitochondrial o2 consumption and atp generation in rat proximal tubule cells. American Journal of Physiology-Renal Physiology 0(0): null, 2019. PMID: 31790302.

[18] Edwards Aurélie, Castrop Hayo, Laghmani Kamel, Vallon Volker, and Layton Anita T. Effects of nkcc2 isoform regulation on nacl transport in thick ascending limb and macula densa: a modeling study. American Journal of Physiology-Renal Physiology 307(2): F137–F146, 2014.

[19] Evans Roger G, Goddard Duncan, Eppel Gabriela A, and O’Connor Paul M. Factors that render the kidney susceptible to tissue hypoxia in hypoxemia. American Journal of Physiology-Regulatory, Integrative and Comparative Physiology 300(4): R931–R940, 2011.

[20] Feldkamp Thorsten, Weinberg Joel M, Hörbelt Markus, Von Kropff Christina, Witzke Oliver, Nürnberger Jens, and Kribben Andreas. Evidence for involvement of nonesterified fatty acid-induced protonophoric uncoupling during mitochondrial dysfunction caused by hypoxia and reoxygenation. Nephrology Dialysis Transplantation 24(1): 43–51, 2009.

[21] Forbes Josephine M and Thorburn David R. Mitochondrial dysfunction in diabetic kidney disease. Nature Reviews Nephrology 14(5): 291, 2018.

[22] Forman NG and Wilson DF. Dependence of mitochondrial oxidative phosphorylation on activity of the adenine nucleotide translocase. Journal of Biological Chemistry 258(14): 8649–8655, 1983.

[23] Formentini Laura, Pereira Marta P, Sánchez-Cenizo Laura, Santacatterina Fulvio, Lucas José J, Navarro Carmen, Martínez-Serrano Alberto, and Cuezva José M. In vivo inhibition of the mitochondrial h+-atp synthase in neurons promotes metabolic preconditioning. The EMBO journal 33(7): 762–778, 2014.

[24] Freeman Dominique M, Bartlett Sylvia, Radda George, and Ross Brian. Energetics of sodium transport in the kidney: saturation transfer 31p-nmr. Biochimica et Biophysica Acta (BBA)-Molecular Cell Research 762(2): 325–336, 1983.

[25] Freeman Dominique M, Chan Laurence, Yahaya Haliru, Holloway Paul, and Ross Brian D. Magnetic resonance spectroscopy for the determination of renal metabolic rate in vivo. Kidney international 30(1): 35–42, 1986.

[26] Friederich-Persson M, Persson Patrik, Hansell Peter, and Palm Fredrik. Deletion of uncoupling protein-2 reduces renal mitochondrial leak respiration, intrarenal hypoxia and proteinuria in a mouse model of type 1 diabetes. Acta Physiologica 223(4): e13058, 2018.

[27] Fry Brendan C, Edwards Aurélie, Sgouralis Ioannis, and Layton Anita T. Impact of renal medullary three-dimensional architecture on oxygen transport. American Journal of Physiology-Renal Physiology 307(3): F263–F272, 2014.

[28] Galgamuwa Ramindhu, Hardy Kristine, Dahlstrom Jane E, Blackburn Anneke C, Wium Elize, Rooke Melissa, Cappello Jean Y, Tummala Padmaja, Patel Hardip R, Chuah Aaron, and others. Dichloroacetate prevents cisplatin-induced nephrotoxicity without compromising cisplatin anticancer properties. Journal of the American Society of Nephrology 27(11): 3331–3344, 2016.

[29] Galgamuwa Ramindhu, Hardy Kristine, Dahlstrom Jane E, Blackburn Anneke C, Wium Elize, Rooke Melissa, Cappello Jean Y, Tummala Padmaja, Patel Hardip R, Chuah Aaron, and others. Dichloroacetate prevents cisplatin-induced nephrotoxicity without compromising cisplatin anticancer properties. Journal of the American Society of Nephrology 27(11): 3331–3344, 2016.

[30] Gershon Nahum D, Porter Keith R, and Trus Benes L. The cytoplasmic matrix: its volume and surface area and the diffusion of molecules through it. Proceedings of the National Academy of Sciences 82(15): 5030–5034, 1985.

[31] Hall Andrew M, Bass Paul, and Unwin Robert J. Drug-induced renal fanconi syndrome. QJM: An International Journal of Medicine 107(4): 261–269, 2014.

[32] Hall Andrew M, Rhodes George J, Sandoval Ruben M, Corridon Peter R, and Molitoris Bruce A. In vivo multiphoton imaging of mitochondrial structure and function during acute kidney injury. Kidney international 83(1): 72–83, 2013.

[33] Hall Andrew M and Unwin Robert J. The not so ‘mighty chondrion’: emergence of renal diseases due to mitochondrial dysfunction. Nephron Physiology 105(1): p1–p10, 2007.

[34] Hems DA and Gaja G. Carbohydrate metabolism in the isolated perfused rat kidney. Biochemical Journal 128(2): 421–426, 1972.

[35] Hoffman David L, Salter Jason D, and Brookes Paul S. Response of mitochondrial reactive oxygen species generation to steady-state oxygen tension: implications for hypoxic cell signaling. American Journal of Physiology-Heart and Circulatory Physiology 292(1): H101–H108, 2007.

[36] Hu Rui and Layton Anita. A computational model of kidney function in a patient with diabetes. International Journal of Molecular Sciences 22(11): 5819, 2021.

[37] Hu Rui, McDonough Alicia A, and Layton Anita T. Functional implications of the sex differences in transporter abundance along the rat nephron: modeling and analysis. American Journal of Physiology-Renal Physiology 317(6): F1462–F1474, 2019.

[38] Hu Rui, McDonough Alicia A, and Layton Anita T. Sex differences in solute transport along the nephrons: effects of na+ transport inhibition. American Journal of Physiology-Renal Physiology 319(3): F487–F505, 2020.

[39] Jacobs Howard T. The mitochondrial theory of aging: dead or alive? Aging cell 2(1): 11–17, 2003.

[40] Klastersky Jean. Side effects of ifosfamide. Oncology 65(Suppl. 2): 7–10, 2003.

[41] Komlódi Tımea, Geibl Fanni F, Sassani Matilde, Ambrus Attila, and Tretter László. Membrane potential and delta ph dependency of reverse electron transport-associated hydrogen peroxide production in brain and heart mitochondria. Journal of bioenergetics and biomembranes 50(5): 355–365, 2018.

[42] Krebs Hans Adolf and Johnson William Arthur. Metabolism of ketonic acids in animal tissues. Biochemical Journal 31(4): 645, 1937.

[43] Lambert Adrian J and Brand Martin D. Superoxide production by nadh: ubiquinone oxidoreductase (complex i) depends on the ph gradient across the mitochondrial inner membrane. Biochemical Journal 382(2): 511–517, 2004.

[44] Layton Anita T, Edwards Aurélie, and Vallon Volker. Adaptive changes in gfr, tubular morphology, and transport in subtotal nephrectomized kidneys: modeling and analysis. American Journal of Physiology-Renal Physiology 313(2): F199–F209, 2017.

[45] Layton Anita T, Edwards Auŕelie, and Vallon Volker. Renal potassium handling in rats with subtotal nephrectomy: modeling and analysis. American Journal of Physiology-Renal Physiology 314(4): F643–F657, 2018.

[46] Layton Anita T, Laghmani Kamel, Vallon Volker, and Edwards Aurélie. Solute transport and oxygen consumption along the nephrons: effects of na+ transport inhibitors. American Journal of Physiology-Renal Physiology 311(6): F1217–F1229, 2016.

[47] Layton Anita T and Vallon Volker. Sglt2 inhibition in a kidney with reduced nephron number: modeling and analysis of solute transport and metabolism. American Journal of Physiology-Renal Physiology 314(5): F969–F984, 2018.

[48] Layton Anita T, Vallon Volker, and Edwards Aurélie. Modeling oxygen consumption in the proximal tubule: effects of nhe and sglt2 inhibition. American Journal of Physiology-Renal Physiology 308(12): F1343–F1357, 2015.

[49] Layton Anita T, Vallon Volker, and Edwards Aurélie. A computational model for simulating solute transport and oxygen consumption along the nephrons. American Journal of Physiology-Renal Physiology 311(6): F1378–F1390, 2016.

[50] Layton Anita T, Vallon Volker, and Edwards Aurélie. Predicted consequences of diabetes and sglt inhibition on transport and oxygen consumption along a rat nephron. American Journal of Physiology-Renal Physiology 310(11): F1269–F1283, 2016.

[51] Layton AT and Edwards A. Mathematical Modeling in Renal Physiology, volume 862. Springer, 2014.

[52] LeFurgey Ann, Spencer Anthony J, Jacobs William R, Ingram Peter, and Mandel Lazaro J. Elemental microanalysis of organelles in proximal tubules. i. alterations in transport and metabolism. Journal of the American Society of Nephrology 1(12): 1305– 1320, 1991.

[53] Legrand Matthieu, Mik Egbert G, Johannes Tanja, Payen Didier, and Ince Can. Renal hypoxia and dysoxia after reperfusion of the ischemic kidney. Molecular medicine 14(7): 502–516, 2008.

[54] Lemieux Guy, Berkofsky James, and Lemieux Christiane. Renal tissue metabolism in the rat during chronic metabolic alkalosis: importance of glycolysis. Canadian journal of physiology and pharmacology 64(11): 1419–1426, 1986.

[55] Letellier T, Bédes F, Malgat M, Korzeniewski B, Jouaville LS, and Morkuniene Ramuné. Metabolic control analysis and threshold effect in oxidative phosphorylation: implications for mitochondrial pathologies. Molecular and cellular biochemistry 174(1): 143–148, 1997.

[56] Lewis William, Day Brian J, and Copeland William C. Mitochondrial toxicity of nrti antiviral drugs: an integrated cellular perspective. Nature Reviews Drug Discovery 2(10): 812–822, 2003.

[57] Li Qianyi, McDonough Alicia A, Layton Harold E, and Layton Anita T. Functional implications of sexual dimorphism of transporter patterns along the rat proximal tubule: modeling and analysis. American Journal of Physiology-Renal Physiology 315(3): F692– F700, 2018.

[58] Lieberthal Wilfred, Menza Sarah A, and Levine Jerrold S. Graded atp depletion can cause necrosis or apoptosis of cultured mouse proximal tubular cells. American Journal of Physiology-Renal Physiology 274(2): F315–F327, 1998.

[59] Ma Shuai, Evans Roger G, Iguchi Naoya, Tare Marianne, Parkington Helena C, Bellomo Rinaldo, May Clive N, and Lankadeva Yugeesh R. Sepsis-induced acute kidney injury: A disease of the microcirculation. Microcirculation 26(2): e12483, 2019.

[60] Miyahara JT and Karler R. Effect of salicylate on oxidative phosphorylation and respiration of mitochondrial fragments. Biochemical Journal 97(1): 194–198, 1965.

[61] Munusamy Shankar and MacMillan-Crow Lee Ann. Mitochondrial superoxide plays a crucial role in the development of mitochondrial dysfunction during high glucose exposure in rat renal proximal tubular cells. Free Radical Biology and Medicine 46(8): 1149–1157, 2009.

[62] Nissim Itzhak, Horyn Oksana, Daikhin Yevgeny, Nissim Ilana, Luhovyy Bohdan, Phillips Peter C, and Yudkoff Marc. Ifosfamide-induced nephrotoxicity: mechanism and prevention. Cancer research 66(15): 7824–7831, 2006.

[63] O’Reilly Molly, Young Luke, Kirkwood Nerissa K, Richardson Guy P, Kros Corné J, and Moore Anthony L. Gentamicin affects the bioenergetics of isolated mitochondria and collapses the mitochondrial membrane potential in cochlear sensory hair cells. Frontiers in cellular neuroscience 13: 416, 2019.

[64] Pfaller Walter and Rittinger Michael. In: Biochemical Aspects of Renal Function. Elsevier, 1980, p. 17–22.

[65] Purvis JL and Slater EC. The effect of magnesium on oxidative phosphorylation and mitochondrial adenosine triphosphatase. Experimental Cell Research 16(1): 109–117, 1959.

[66] Quiros Yaremi, Vicente-Vicente Laura, Morales Ana I, López-Novoa José M, and López-Hernández Francisco J. An integrative overview on the mechanisms underlying the renal tubular cytotoxicity of gentamicin. Toxicological sciences 119(2): 245–256, 2011.

[67] Schaefer Andrew M, Taylor Robert W, Turnbull Douglass M, and Chinnery Patrick F. The epidemiology of mitochondrial disorders—past, present and future. Biochimica et Biophysica Acta (BBA)-Bioenergetics 1659(2-3): 115–120, 2004.

[68] Schiffer Tomas A, Christensen Michael, Gustafsson Håkan, and Palm Fredrik. The effect of inactin on kidney mitochondrial function and production of reactive oxygen species. PloS one 13(11): e0207728, 2018.

[69] Schiffer Tomas A, Gustafsson Håkan, and Palm Fredrik. Kidney outer medulla mitochondria are more efficient compared with cortex mitochondria as a strategy to sustain atp production in a suboptimal environment. American Journal of Physiology-Renal Physiology 315(3): F677–F681, 2018.

[70] Schirris Tom JJ, Renkema G Herma, Ritschel Tina, Voermans Nicol C, Bilos Albert, Engelen van Baziel GM, Brandt Ulrich, Koopman Werner JH, Beyrath Julien D, Rodenburg Richard J, and others. Statin-induced myopathy is associated with mitochondrial complex iii inhibition. Cell metabolism 22(3): 399–407, 2015.

[71] Schmitt Sabine, Schulz Sabine, Schropp Eva-Maria, Eberhagen Carola, Simmons Alisha, Beisker Wolfgang, Aichler Michaela, and Zischka Hans. Why to compare absolute numbers of mitochondria. Mitochondrion 19: 113–123, 2014.

[72] Schwerdt Gerald, Gordjani Nader, Benesic Andreas, Freudinger Ruth, Wollny Brigitte, Kirchhoff Antje, and Gekle Michael. Chloroacetaldehyde-and acrolein-induced death of human proximal tubule cells. Pediatric nephrology 21(1): 60–67, 2006.

[73] Secker Philipp F, Beneke Sascha, Schlichenmaier Nadja, Delp Johannes, Gutbier Simon, Leist Marcel, and Dietrich Daniel R. Canagliflozin mediated dual inhibition of mitochondrial glutamate dehydrogenase and complex i: an off-target adverse effect. Cell death & disease 9(2): 1–13, 2018.

[74] Sgouralis Ioannis, Evans Roger G, Gardiner Bruce S, Smith Julian A, Fry Brendan C, and Layton Anita T. Renal hemodynamics, function, and oxygenation during cardiac surgery performed on cardiopulmonary bypass: a modeling study. Physiological reports 3(1): e12260, 2015.

[75] Sgouralis Ioannis, Evans Roger G, and Layton Anita T. Renal medullary and urinary oxygen tension during cardiopulmonary bypass in the rat. Mathematical medicine and biology: a journal of the IMA 34(3): 313–333, 2017.

[76] Somlyo Andrew P, Somlyo Avril V, and Shuman Henry. Electron probe analysis of vascular smooth muscle. composition of mitochondria, nuclei, and cytoplasm. The Journal of cell biology 81(2): 316–335, 1979.

[77] Stacpoole Peter W. Therapeutic targeting of the pyruvate dehydrogenase complex/pyruvate dehydrogenase kinase (pdc/pdk) axis in cancer. JNCI: Journal of the National Cancer Institute 109(11), 2017.

[78] Stillman Isaac E, Brezis Mayer, Heyman Samuel N, Epstein Franklin H, Spokes Kate, and Rosen Seymour. Effects of salt depletion on the kidney: changes in medullary oxygenation and thick ascending limb size. Journal of the American Society of Nephrology 4(8): 1538–1545, 1994.

[79] Suen Der-Fen, Norris Kristi L, and Youle Richard J. Mitochondrial dynamics and apoptosis. Genes & development 22(12): 1577–1590, 2008.

[80] Sun SQ, Shi S-L, Hunt JA, and Leapman RD. Quantitative water mapping of cryosectioned cells by electron energy-loss spectroscopy. Journal of microscopy 177(1): 18–30, 1995.

[81] Tapiawala Shruti N, Tinckam Kathryn J, Cardella Carl J, Schiff Jeffrey, Cattran Daniel C, Cole Edward H, and Kim S Joseph. Delayed graft function and the risk for death with a functioning graft. Journal of the American Society of Nephrology 21(1): 153–161, 2010.

[82] Tsimaratos M, Roger F, Chabardes D, Mordasini D, Hasler U, Doucet A, Martin P-Y, and Feraille E. C-peptide stimulates na+, k+-atpase activity via pkc alpha in rat medullary thick ascending limb. Diabetologia 46(1): 124–131, 2003.

[83] Vrbjar N, Javorkova V, and Pechanova O. Changes of sodium and atp affinities of the renal na, k-atpase during and after nitric oxide-deficient hypertension. Physiological research 51(5): 475–482, 2002.

[84] Weinberg Joel M and Humes H David. Mechanisms of gentamicin-induced dysfunction of renal cortical mitochondria: I. effects on mitochondrial respiration. Archives of biochemistry and biophysics 205(1): 222–231, 1980.

[85] Wu Fan, Yang Feng, Vinnakota Kalyan C, and Beard Daniel A. Computer modeling of mitochondrial tricarboxylic acid cycle, oxidative phosphorylation, metabolite transport, and electrophysiology. Journal of Biological Chemistry 282(34): 24525–24537, 2007.

[86] Yu Hai-Tao, Fu Xiao-Yi, Liang Bing, Wang Shuang, Liu Jian-Kang, Wang Shu-Ran, and Feng Zhi-Hui. Oxidative damage of mitochondrial respiratory chain in different organs of a rat model of diet-induced obesity. European journal of nutrition 57(5): 1957–1967, 2018.

[87] Zager Richard A. Mitochondrial free radical production induces lipid peroxidation during myohemoglobinuria. Kidney international 49(3): 741–751, 1996.

[88] Zhang Wensheng and Edwards Aurélie. Oxygen transport across vasa recta in the renal medulla. American Journal of Physiology-Heart and Circulatory Physiology 283(3): H1042–H1055, 2002.

[89] Zoratti M, Favaron M, Pietrobon D, and Petronilli V. Nigericin-induced transient changes in rat-liver mitochondria. Biochimica et Biophysica Acta (BBA)-Bioenergetics 767(2): 231–239, 1984.

